# Evolutionary unpredictability in cancer model system

**DOI:** 10.1101/2022.06.01.494285

**Authors:** Subhayan Chattopadhyay, Jenny Karlsson, Adriana Mañas, Ryu Kanzaki, Elina Fredlund, Andrew J. Murphy, Christopher L. Morton, Natalie Andersson, Mary A. Woolard, Karin Hansson, Katarzyna Radke, Andrew M. Davidhoff, Sofie Mohlin, Kristian Pietras, Daniel Bexell, David Gisselsson

**Author notes:** Corresponding Author: Subhayan Chattopadhyay, Division of Clinical Genetics, Department of Laboratory Medicine, Lund University, SE22184 Lund, Sweden.

## Abstract

Despite the advent of personalized medicine, it is still difficult to predict how a cancer develops over time at the level of the individual patient or even in cancer model systems which begs the question whether certain aspects of cancer can ever be predicted or if there is an inherent unpredictability in cancer, similar to other complex biological systems, We demonstrate by a combination of agent-based mathematical modelling and analysis of data from patient-derived xenograft systems from multiple cancer types that certain conditions may invoke chaotic fluctuations in the clonal landscape of cancer cells. Our findings indicate that under those conditions, the cancer genome behaves as a complex dynamic system, making its long-term evolution inherently unpredictable.

## Introduction

A quarter of a century ago, in 1993, Donald S. Coffey hypothesized that cancer is an ‘abrupt’ and ‘emergent’ phenomenon caused by the transformation of the cell proliferation machinery from an ordered to a disordered albeit self-organizing state ^1^. In the following decades, researchers have increasingly focused on the characterization of dysregulated (disordered) genomic and non-genomic elements in cancer. However, the prevalent somatic mutation theory stating that near-random DNA lesions combined with disease-specific selection cause cancer still indicates a failure to grasp Coffey’s vision of using the self-organizing features of cancer to identify unifying aspects across malignancies ^2-4^. With the advent of precision cancer medicine, more efforts are now being made to characterize the broad spectrum of genetic variation among individual tumors, instead of describing unifying features among cancers. This has resulted in a fine-tuned prognostication of many neoplasms and the identification of treatment targets based on the molecular signatures of each cancer. However, in many cases, the remaining threat of a relapse at an unknown time point, often manifesting as treatment-resistant aggressive disease, reminds us that cancer shares features of resilience with many other self-organizing systems. Cancer’s relapse mechanisms have been thoroughly studied through the lens of genetic diversity, elucidating how tumors evolve along different evolutionary trajectories ^5^ and how resistant clones often appear due to excessive evolutionary branching early in disease, leading to dormant tumor cell populations with a potential to clonally fixate afterwards ^6-7^. Despite the massive amount of data accumulated on the molecular routes of relapse, it remains an essentially unpredictable phenomenon. However, as inferred by Coffey, cancer is not a stochastic phenomenon. Instead, it results from runaway dysregulation in a complex, dynamic, and adaptive system. Can we use generic knowledge from other systems in a state near chaos to better understand tumorigenesis?

Today, the generalized logistic model (with the Gompertz curve being a special case) is a commonly used construct for simulating spatially constricted (bonded) growth of species/cell populations, and it is popularly used to emulate solid tumor growth ^8-10^. Several comparative studies analyzing *in-vivo* tumor growth have established the suitability of logistic functions in tumor growth estimation ^10-11^. Additionally, it has been used to illustrate clonal selection and genetic drift in solid tumors in silico ^12^. Simulations have shown that the evolutionary trajectories of cancers are highly dependent on how cell populations grow and how they interact with the stromal boundary ^13^. However, whether the logistic function as a mechanism for tumor growth can reliably predict emergent clonal geographies in tumorigenesis remains to be explored. Here, we probe this question with a focus on the predictability of mutational landscapes.

## Results and Discussion

First, we performed simulations based on the assumption that tumors, prior to clonal expansion, emerge from a uniform population of cells that are henceforth referred to as ancestors. Ancestors underwent clonal expansion adhering to certain parameters (i.e., a growth rate governed by birth and death rates, the rate of acquiring mutations at each cell division, and the probability of a mutation to be a driver mutation), which were set at initiation and remained unchanged, purely for the sake of simplicity. The selective advantages provided by a mutation were sourced from the deleteriousness scores provided by the COSMIC database (detailed in Methods) ^14^ and the maximum number of cells at the end of the simulation was kept fixed. To monitor clonal landscapes at the end of simulations, we focused on how the cells underwent genetic diversification due to variations in simulation parameters, making them genetically distant and heterogeneous progenies of the ancestors. All the cells at the end of each simulation were clustered according to the number of acquired mutations, and the least mutated cluster was considered to represent the most recent common ancestors. With this, we arrive at a simple measure of the percentage of ancestors remaining at the end of the simulation run.

In his seminal work on chaotic oscillations in dynamic systems, James A. Yorke showed that a self-repeating process with a periodicity of three or more experiences chaotic fluctuations. This was exemplified by the logistic function ^15^, which is a sigmoidal curve with a periodically oscillating slope (fig. 1A). Robert May further demonstrated how chaotic fluctuations could appear as a function of logistic growth in a bifurcation diagram ^16^ (fig. 1B), presenting how a function can assume more than one value near its asymptotes (orbits). Next, we evaluated the change in the percentage of the remaining ancestors against varying initial conditions to observe possible indications of dynamic oscillation.

**Figure 1.**
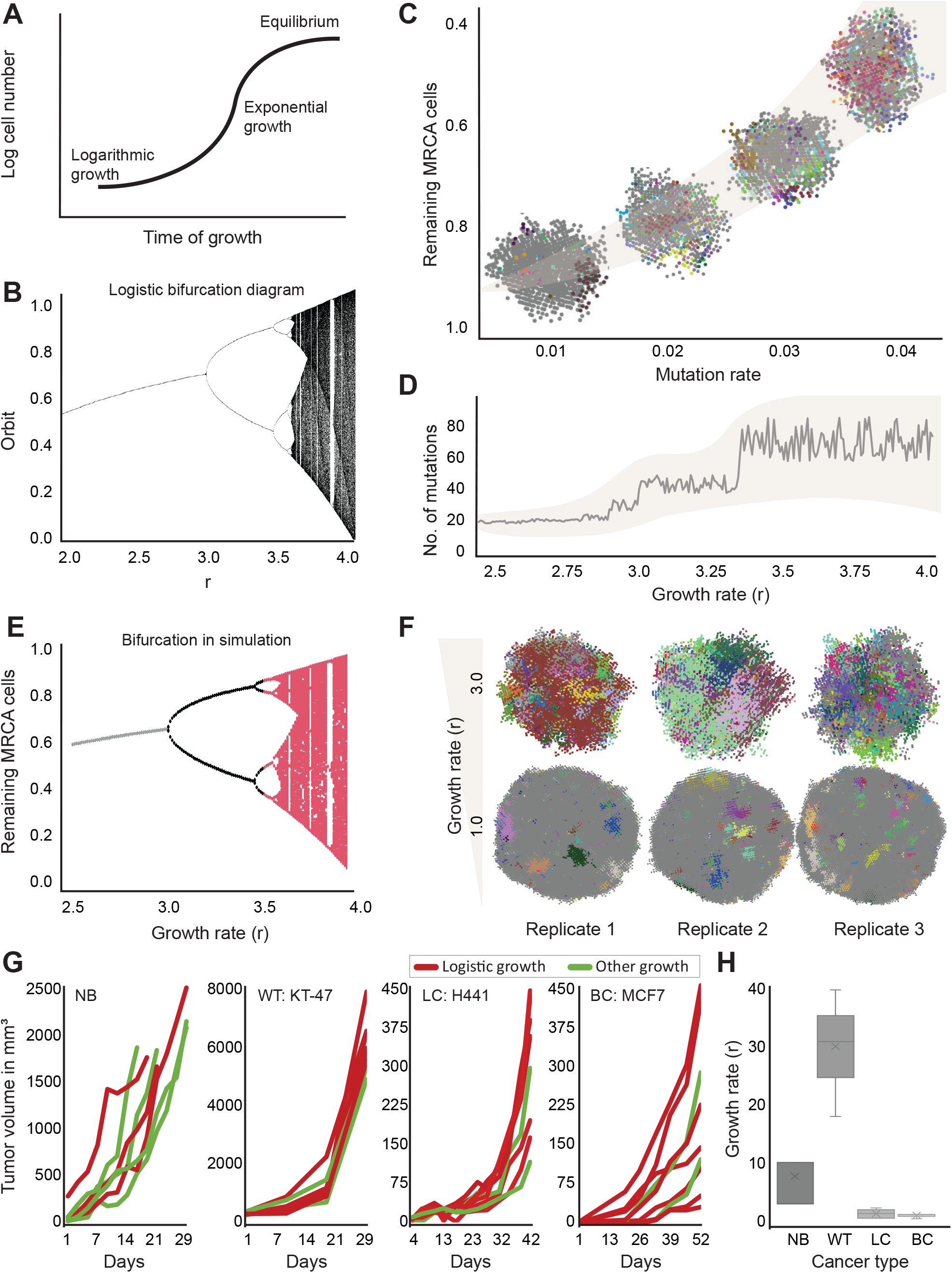
Simulation of tumor evolution. (**A**) Interpretation of the classical shape of a logistic growth curve. (**B**) Classical orbit diagram of logistic curve (logistic map) ^16^. (**C**) Change in percentage of remaining MRCA (most recent common ancestors) cells was plotted against increasing mutation rate. Examples were overlayed at the estimated mean calculated with 100 simulation runs. Confidence interval with shaded overlay depicts 3σ limit. (**D**) Average number of total distinct mutations at the end of simulation is plotted against growth rates (r). All estimates are calculated over 100 separate runs. As logistic map experiences chaotic fluctuations starting at r = 3.0, the x-axis is terminated at 4.0 as it already provided adequate range (confidence interval depicts 3σ limit) (**E**) Cell growth rate was plotted against percentage of remaining ancestors at the end of simulations, grey section indicates r < 3.0, calculated by taking average; black section indicates 3.0 < r < 3.5, calculated by cluster centroids; red section indicates r > 3.5 where all estimates are plotted. (**F**) Three replicates of simulation are shown for growth rates (r) 1.0 and 3.0 to draw attention to the fact that high r results in markedly different mutational pattern compared to that in low r. The colors in each simulation indicate a cluster of clonal cells. (**G**) Tumor growth data in mouse models are depicted; red: logistic growth and green: did not fit to logistic function. NB: neuroblastoma (Radke et. al.); WT: Wilms tumor (KT-47 sample, Murphy et. al.); LC: lung cancer (H441 cell line); BC: breast cancer (MCF7 cell line). (**H**) Box plots of growth rates of data shown in (1-G) plotted across cancer types.

With each simulation cycle, mutations were drawn at random from a set of known somatic mutations (see Methods). Using fixed initial conditions, we evaluated whether genetic diversification remained somewhat predictable, given certain mutation parameters. Keeping the growth rate unchanged, we first varied the probability of acquiring a mutation at each cell division between 0.01 and 0.04 (fig. 1C) ^12^. Cellular growth rates were affected by mutation rate, mutation type and mutation aggregation all of which conferred changes in cellular fitness. We also observed a nonlinear monotonically decreasing relationship between the percentage of ancestors and increasing mutation rate. The fraction of the remaining ancestors at the end of the simulation varied on average between 92% (low mutation rate) and 36% (high mutation rate, calculated over 100 simulation runs). However, the relationship between the median number of mutations and growth rate was less straightforward. Mutation acquisition seemed to undergo a dramatic step-like increment over an arbitrary span (fig. 1D). Clear jumps in median mutation aggregation was observed near growth rates 3.0 and 3.4, indicating rapid changes in the distribution of mutation aggregation at certain growth rates. Indeed, the jumps in mutation aggregation corresponded to the classic bifurcation diagram of the logistic function (fig. 1B ^16^), when plotting the percentage of the remaining ancestor against the growth rate (fig. 1E). A faster growth rate resulted in a markedly heterogeneous mutational landscape compared to a slow growth rate (fig. 1F). In principle, the shape of the bifurcation diagram should predict chaotic behavior as the growth constant increases over a certain threshold that also represents a one-to-many solution of the logistic map at the asymptotes. In our simulations, this was observed at a growth rate above 3.0. This led us to conclude that chaotic growth (at least in our case) is a biologically emergent feature that can occur in tumors following logistic growth above a certain rate.

Talkington and Durett evaluated *in vivo* growth characteristics of several cancer cell lines and found that numerous cell lines at least partly resemble logistic (specifically, Gompertzian) growth ^11^ Some of the earliest investigations on *in vitro* growth experiments have also pointed towards a similar pattern ^17-18^. However, the extent to which fast logistic growth occurs in tumors *in vivo* remains unclear.

To evaluate whether fast-growing tumors with potential non-linear clonal evolution exist, we assessed how often logistic growth is observed across *in vivo* model systems in situations simulating relapse by implantation of a limited tumor population. Pediatric cancers are well known to have much higher growth rates than adult cancers. Neuroblastoma (NB) and Wilms tumor (WT) are two of the most common solid pediatric tumors, notorious for their rapid growth. We estimated growth rates from previously published data on untreated patient-derived xenografts (PDX) derived from NB and WT ^5,19-22^. For comparison with adult tumors (i.e., slow growers) ^23^, we also evaluated the uninhibited growth of several lung and breast cancer cell lines (unpublished data). Strikingly, 43% of the evaluated NB PDXs abided by logistic growth (73% of growth rates were more than 3.0, with a median of 10.0), considerably above the chaotic bifurcation limit of 3.0, whereas 75% of the WT PDXs showed logistic growth (all growth rates over 3.0) with a median growth rate of 31.0 (fig. 1G-H). PDXs from H441 lung cancer and MCF7 breast cancer cell lines, 71% and 78%, respectively, experienced logistic growth, but none over 3.0. The median growth rates were only 1.13 and 0.9, respectively, far below that for chaotic fluctuations. In addition to patient-derived models, we also evaluated the growth of two lineages of the NB SK-N-BE(2)C cell line *in vivo*, with median growth rates of 5.0 and 4.0%, respectively (Supplementary material). When combined, the growth rates from all NB replicates had a median of 6.0, and that of the WT was 24.0. All but one breast cancer replicate that adhered to logistic growth had growth rates below 3.0, with a median of 0.99 and that for lung cancer replicates was 0.68, none reaching 3.0.

The implications of the present study are limited to tumors that demonstrated a logistic growth curve; however, not all tumors did so. This is possibly due to different inherent growth characteristics, as the absolute volume of the tumors varies substantially between pediatric and adult tumors. By week four of observation, the average volume for the NB replicates exceeded 2000 mm^3^. In the same time frame adult tumors grew only about 400 mm^3^ except for one replicate (m3) of the MDA-MB-231 breast cancer cell line (growth rate 3.6). The WT PDXs at implantation were all larger than 200 mm^3^ with an average size of 308 mm^3^ which grew to 3800 mm^3^ on average by week 3. Notably, NB PDXs underwent orthotopic implantation initially and were then grafted heterotopically^22^ whereas the WTs were heterotopic. Often, the reason that a curve failed to reasonably fit a logistic function was due to abrupt changes in the growth pattern (measurement artifacts, etc.), which is a regular phenomenon. Nevertheless, the presence of any systematic artifact responsible for the low growth rates in the adult tumors was ruled out, as the experiments yielded growth rate within-variances of 0.023 and 0.012.

Simulations under chaotic growth resulted in massively varied mutational landscapes, implying that evolutionary trajectories, even in controlled model systems, are unpredictable under certain conditions that invoke non-linearity. The concept of genome chaos during cancer evolution has been used to describe non-continuous accumulation of genetic lesions^24^. This model claims that tumor evolution is bi-phasic, starting with punctuated aggregation of mutations followed by a maintenance phase. The phase transition depends on evolutionary stress and also determines the rate at which the tumor grows/evolves. The co-dependence of the phases is argued to result in macroevolutionary chaos originating from a predictable microevolutionary background^25^.Our experimental data imply that non-linear evolution close to chaos is particularly likely to occur in fast-growing (pediatric) tumors. The mouse model data illustrate that some tumor types are associated with specific growth characteristics (non-logistic/logistic and slow/fast growing), which determine whether they evolve clonally in a chaotic or predictable fashion. Despite the correlative evidence, whether a logistic growth rate above the threshold for bifurcating states demonstrably invokes genomic chaos remains to be validated. We simulated the population expansion process with a discrete growth following a logistic equation. This enabled us to implement an analytical approach to spatial simulation, but to some degree restricted genetic evolution as the population numbers reached at each generation were set to follow a logistic curve. Evolutionary dynamics for tumors growing in a non-logistic fashion need also to be compared alongside logistically growing tumors/cell populations to evaluate if chaotic fluctuations are specific to logistic growth. According to the punctuated evolutionary model for cancer^24^, the maintenance phase acts as a self-regulatory mechanism to reorganize cells after a period of arbitrary mutation aggregation. It may be that chaos arises as a result of the dysregulation in these two phases of evolution in a way that voids the natural equilibrium of the evolutionary homeostasis.

Our findings suggest an intrinsic limit on what can be predicted using today’s approach of precision oncology. However, our findings indicate that observing the tumor growth rate could be a potential workaround to determine the degree of predictability the tumor is expected to evolve, in turn providing insights into how often the clonal landscape of a tumor needs to be resampled to evaluate options for targeted therapy based on the molecular profile.

## Methods

### Assumptions underlying *in silico* modelling

Here, we simulated a synthetic model for tumorigenesis, incorporating some of the basic assumptions of the somatic mutation theory. It remains to be seen whether DNA damage-driven neutral or Darwinian evolution becomes apparent with this system, as the model does not assume a position on either dogma of continuous tumor evolution. The model focuses on simulating evolutionary pockets in a tumor soma that vary across its geography, similar to how it is explained by Waclaw and colleagues^12^.

We begin the simulation with a predetermined number of cells, C, indicating a fixed carrying capacity (maximum number of cells in homeostasis) in a three-dimensional automaton where all but a few (randomly chosen, < 10) neighboring cells are considered to be non-neoplastic at the beginning (generation 0). Over the total span of each simulation, we modulated the following five properties and treated them as simulation parameters:

- Cell division rate / birth rate (**b**)
- Cell death rate (**d**)
- Mutation rate (**m**)
- Driver mutation probability (**p**)
- Selective advantage / fitness (**b**)

These parameters in tandem control the size of the tumor, how fast it grows, and how genetically diverse and evolved it is at the end of the simulation. As the automaton simulates cellular interaction and not the expansion magnitude, we iterated the evolutionary simulation over punctuated time points. The population size is obtained from a discrete logistic growth with variable growth rate solved analytically as a function of discrete time. The generalized discrete form is given as follows:

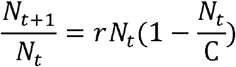

Which can be analytically simplified to the following^26^ where C is carrying capacity of the system, *N*_*j*_ is the number of tumor cells at any given time *j* and *r* is the growth rate.

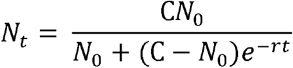

We solved for *N*_*t*_ iteratively for time *t* = 0,1,2, … proxied by the set of whole numbers and simulated the discretized evolutionary dynamic for a tumor expanding from *N*_0_ cells to *N*_*t*_ cells. We implemented these dynamics in the simulation by assuming certain base hypotheses.

#### Conjecture 1

Mutational burden is predicted by the type of mutation and the mutation rate.

At each division any given daughter cell acquires some mutational burden quantified by D_n_; n∈□. The number of neoplastic cells, n, in the automata dictates when the simulation ends, that is, when n = N. Hence, we leverage it as a proportion of time. Since D_n_ changes depending on the mutation rate, and on the effect the accumulated mutations have on the cell, it follows that D_n_ is a function of both **m** and **p**

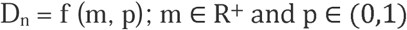

#### Conjecture 2

Cell birth and death rates depend on mutational burden.

Every k^th^ cell in the population undergoes cell division after time T (k_birth, n_) and apoptosis after time T (k_death, n_); n, k ∈ ℤ^+^, k ∈K < N. These are proxies for *birth rate* and *death rate* and, therefore, must functionally depend on the acquired mutational burden.

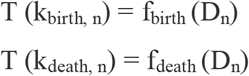

#### Conjecture 3

Acquiring certain mutations creates a positive feedback loop.

With this, we assume that the probability of any cell acquiring mutational burden is predicated by pre-existing lesions in its DNA. At any i^th^ cell division, the probability of a cell acquiring additional mutational burden is dependent on its preexisting burden after the i−1^th^ cell division. Because the cumulative mutational load defines the selective advantage that a cell harbors at any given point, s_i_ = f (s_i-1_), where s_i_ and s_i-1_ are the selective advantages of a cell at each stage and take the functional form of a Markov process, respectively. The selection coefficient is thus shaped indirectly by mutational burden.

### Modeling acquisition of mutations

In this study, we considered only somatic point mutations extracted from publicly available datasets. This is in no way to indicate a lack of availability of data on structural chromosomal changes as well as longer variations, but only due to the unsubstantiated mechanics of conferring selective advantage or disadvantage. We also did not consider inherited mutations, as all cells should harbor the same set of variations, which in turn should have scalable effects across the entire cell population. How these factors affect the evolutionary process is of course of interest, but we assume that they do not affect the relative fitness of the cells. Hence, we adapted a system of evolutionary dynamics that is only affected by point mutations, and to this end, we used their pathogenicity scores.

We extracted the COSMIC v90 dataset (https://cancer.sanger.ac.uk/cosmic/download) for all reported point mutations along with the estimated pathogenicity scores (FATHMM-MKLv2.3)^27^. Every mutation was given a score with a lower limit of 0 and an upper limit of 1 with respect to the corresponding cancer site and histology when applicable. COSMIC identifies strong drivers with score stratification which we use to stratify all possible mutations into to groups, *neutral and pathogenic*. A FATHMM score between 0.5 and 0.7 indicates a ‘driver’ and that above 0.7 indicates a ‘strong driver’. We assumed a mutation to be pathogenic if it had a score > 0.5 and the rest were declared neutral. We extracted the pathogenicity scores of only the pathogenic mutations to compute the fitness advantage. By definition, tumor fitness is indifferent to neutral mutations, nevertheless these were indexed in the genetic background. The overaccumulation of neutral mutations pushed the cell division machinery into arrest above a critical threshold (discussed below) although it did not directly influence cellular fitness.

The fitness landscape was shaped on the basis of two main tenets. First, all site-specific driver mutations reported to date are identified in the literature, along with those that are associated with individual hallmarks of cancer and are extracted for tumor simulation of that site. We retained drivers reported in neuroblastoma as we assumed that the number of driver mutations and their strength of adding to the fitness advantage will determine how the clonal geography takes shape. Once such a mutation is acquired by a cell, it confers a selective advantage to cell survival according to its pathogenicity score. Second, neutral mutations confer mutational burden only via accumulation such that their cumulative aggregation over generations triggers apoptosis abiding Muller’s ratchet^28^.

### How neutral mutations affect growth

Muller’s ratchet predicts that the number of neutral mutations will increase over generations. Hence, the carrying capacity of the population with a given number of mutations is affected, which is given by the equation

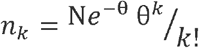

where θ = ^*s*^/_*m*_. Here, n_k_ is the carrying capacity of the cell population harboring neutral mutations. where N is the total population size, m is the mutation rate, and s is the corresponding selection coefficient/fitness. Hence, it can be concluded that the population size-scaled carrying capacity follows a Poisson distribution:

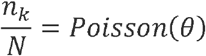

over the random variable K assuming values k∈N^+^ and the carrying capacity of the fittest class of cells, that is, at the beginning of simulation is given by

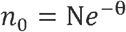

In our simulations, we did not check for sample sizes using a power analysis because our study does not report statistics on between groups or within group variables. No technical replication was completed because the summaries were extracted over multiple runs of simulations.

The simulations were performed with SITH^12^; prediction, fitting and evaluation of the models were performed in R version 4.1.

## Data availability

The script for the simulated data is available on GitHub. All PDX data were published or available from the respective corresponding authors. A summary of the data is provided in the Supplementary Information.

## Ethics statement

Ethical permits for the published results are available in the respective publications.

