## Supplementary for "Evolutionary unpredictability in cancer model system"

**Supplementary Table 1.** Summary of accumulated experiments

| **Author identifier** | **Cancer type** | **sample type** | **No. of replicates** | **Total no. of mice** | **Source** |
|---|---|---|---|---|---|
| Hansson et. al. | Neuroblastoma | PDX | 2 | 11 | https://doi.org/10.1126/scitranslmed.aba4434 |
| Radke et. al. | Neuroblastoma | PDX | 1 | 7 | https://doi.org/10.1016/j.tranon.2021.101149 |
| Mohlin et. al. | Neuroblastoma | cell lines | 5 | 32 | https://doi.org/10.1158/0008-5472.CAN-15-0708; (some unpublished) |
| Manas et. al. | Neuroblastoma | PDX | 6 | 30 | https://doi.org/10.1101/2022.04.01.486670 |
| Murphy et. al. | Wilms tumor | PDX | 6 | 48 | https://doi.org/10.1038/s41467-019-13646-9 |
| n.a. | Breast cancer | cell lines | 2 | 18 | Unpublished data |
| n.a. | Lung cancer | cell lines | 5 | 34 | Unpublished data |

**Supplementary Table 2.** Neuroblastoma PDXs (Hansson et. al.)

Replicate 1.

| Control | Growth rate | %log | %chaos |
|---|---|---|---|
| **A1** | 2.9 | 75% | 67% |
| **A2** | 6.9 | | |
| **A3** | 7.1 | | |
| **A4** | n.a | | |

Replicate 2.

| Control | Growth rate | %log | %chaos |
|---|---|---|---|
| **A1** | 4.1 | 43% | 100% |
| **A2** | n.a | | |
| **A3** | n.a | | |
| **A4** | 4.4 | | |
| **A5** | n.a | | |
| **B1** | 4.9 | | |
| **B2** | n.a | | |

%log: Percentage of mice that followed a logistic growth
%chaos: Percentage of mice following logistic growth that observed growth rate over 3.0

**Supplementary Table 3.** Neuroblastoma PDXs (Radke et. al.)

| Control | Growth rate | %log | %chaos |
|---|---|---|---|
| **A1** | n.a | 43% | 67% |
| **A2** | 2.5 | | |
| **A3** | n.a | | |
| **A4** | 9.8 | | |
| **A5** | n.a | | |
| **A6** | n.a | | |
| **A7** | 8.8 | | |

%log: Percentage of mice that followed a logistic growth
%chaos: Percentage of mice following logistic growth that observed growth rate over 3.0

**Supplementary Table 4.** Neuroblastoma SK-N-BE2(C) cell line (Mohlin et. al.)

Replicate 1.

| Control | Growth rate | %log | %chaos |
|---|---|---|---|
| **A31** | 4.8 | 50% | 80% |
| **A41** | 6.2 | | |
| **B11** | n.a | | |
| **B21** | n.a | | |
| **B31** | 5.8 | | |
| **A32** | 1.7 | | |
| **A42** | 11 | | |
| **B12** | n.a | | |
| **B22** | n.a | | |
| **B32** | n.a | | |

Replicate 2.

| Control | Growth rate | %log | %chaos |
|---|---|---|---|
| **A1** | 1.4 | 60% | 67% |
| **A2** | 3.7 | | |
| **A3** | 4 | | |
| **A4** | n.a | | |
| **A5** | n.a | | |

Replicate 3.

| Control | Growth rate | %log | %chaos |
|---|---|---|---|
| **m1** | n.a | 60% | 100% |
| **m2** | 6.5 | | |
| **m3** | 20 | | |
| **m4** | n.a | | |
| **m5** | 15 | | |

Replicate 4.

| Control | Growth rate | %log | %chaos |
|---|---|---|---|
| **m1** | 5 | 44% | 75% |
| **m2** | n.a | | |
| **m3** | n.a | | |
| **m4** | n.a | | |
| **m5** | n.a | | |
| **m6** | n.a | | |
| **m7** | 1.5 | | |
| **m8** | 5.6 | | |
| **m9** | 3.7 | | |

Replicate 5.

| Control | Growth rate | %log | %chaos |
|---|---|---|---|
| **m1** | 3.9 | 67% | 100% |
| **m2** | n.a | | |
| **m4** | 4.1 | | |

%log: Percentage of mice that followed a logistic growth
%chaos: Percentage of mice following logistic growth that observed growth rate over 3.0

**Supplementary Table 5.** Neuroblastoma PDXs (Manas et. al.)

Model: PDX1 (nude mice)

| Control | Growth rate | %log | %chaos |
|---|---|---|---|
| **c1** | 4.5 | 75% | 100% |
| **c2** | 3.1 | | |
| **c3** | n.a | | |
| **c4** | 11 | | |

Model: PDX1 (NSG mice)

| Control | Growth rate | %log | %chaos |
|---|---|---|---|
| **c1** | 25 | 60% | 100% |
| **c2** | 29 | | |
| **c3** | n.a | | |
| **c4** | n.a | | |
| **c5** | 6 | | |

Model: PDX2 (NSG mice)

| Control | Growth rate | %log | %chaos |
|---|---|---|---|
| **c1** | 4.4 | 71% | 100% |
| **c2** | 13 | | |
| **c3** | 11 | | |
| **c4** | n.a | | |
| **c5** | 16 | | |
| **c6** | 12 | | |
| **c7** | n.a | | |

Model: PDX3 (nude mice), replicate 1

| Control | Growth rate | %log | %chaos |
|---|---|---|---|
| **C1** | 1.8 | 50% | 50% |
| **C2** | 6.4 | | |
| **C3** | n.a | | |
| **C4** | n.a | | |

Model: PDX3 (nude mice), replicate 2

| Control | Growth rate | %log | %chaos |
|---|---|---|---|
| **c1** | n.a | 40% | 100% |
| **c2** | 9.4 | | |
| **c3** | n.a | | |
| **c4** | 9.8 | | |
| **c5** | n.a | | |

Model: PDX3 (nude mice), replicate 3

| Control | Growth rate | %log | %chaos |
|---|---|---|---|
| **c1** | 17 | 80% | 100% |
| **c2** | 18 | | |
| **c3** | n.a | | |
| **c4** | 16 | | |
| **c5** | 16 | | |

%log: Percentage of mice that followed a logistic growth
%chaos: Percentage of mice following logistic growth that observed growth rate over 3.0

**Supplementary Table 6.** Wilms tumor PDXs (Murphy et. al.)

Model: KT47

| Control | Growth rate | %log | %chaos |
|---|---|---|---|
| **m1** | n.a | 75% | 100% |
| **m2** | 30 | | |
| **m3** | 18 | | |
| **m4** | 34 | | |
| **m5** | 32 | | |
| **m6** | 40 | | |
| **m7** | n.a | | |
| **m8** | 27 | | |

Model: KT53

| Control | Growth rate | %log | %chaos |
|---|---|---|---|
| **m1** | 31 | 100% | 100% |
| **m2** | 35 | | |
| **m3** | 37 | | |
| **m4** | 34 | | |
| **m5** | 32 | | |
| **m6** | 32 | | |
| **m7** | 27 | | |
| **m8** | 32 | | |

Model: KT51

| Control | Growth rate | %log | %chaos |
|---|---|---|---|
| **m1** | 21 | 88% | 100% |
| **m2** | 6.8 | | |
| **m3** | 15 | | |
| **m4** | 15 | | |
| **m5** | 40 | | |
| **m6** | n.a | | |
| **m7** | 39 | | |
| **m8** | 15 | | |

Model: KT45

| Control | Growth rate | %log | %chaos |
|---|---|---|---|
| **m1** | n.a | 63% | 100% |
| **m2** | n.a | | |
| **m3** | 5.8 | | |
| **m4** | 20 | | |
| **m5** | 12 | | |
| **m6** | 4.3 | | |
| **m7** | 7.3 | | |
| **m8** | n.a | | |

%log: Percentage of mice that followed a logistic growth
%chaos: Percentage of mice following logistic growth that observed growth rate over 3.0

Model: KT75

| Control | Growth rate | %log | %chaos |
|---|---|---|---|
| **m1** | n.a | 38% | 100% |
| **m2** | n.a | | |
| **m3** | 8.2 | | |
| **m4** | 4.8 | | |
| **m5** | n.a | | |
| **m6** | n.a | | |
| **m7** | n.a | | |
| **m8** | 7.6 | | |

Model: KT43

| Control | Growth rate | %log | %chaos |
|---|---|---|---|
| **m1** | n.a | 13% | 100% |
| **m2** | n.a | | |
| **m3** | 4.4 | | |
| **m4** | n.a | | |
| **m5** | n.a | | |
| **m6** | n.a | | |
| **m7** | n.a | | |
| **m8** | n.a | | |

%log: Percentage of mice that followed a logistic growth
%chaos: Percentage of mice following logistic growth that observed growth rate over 3.0

**Supplementary Table 7.** Breast cancer cell line models

Cell line: MCF7

| Control | Growth rate | %log | %chaos |
|---|---|---|---|
| **m1** | 0.9 | 89% | 0% |
| **m2** | 0.23 | | |
| **m3** | 0.98 | | |
| **m4** | 0.63 | | |
| **m6** | n.a | | |
| **m7** | 0.97 | | |
| **m8** | 0.99 | | |
| **m9** | 0.98 | | |
| **m10** | 0.87 | | |

Cell line: MDA-MB-231

| Control | Growth rate | %log | %chaos |
|---|---|---|---|
| **m1** | 1.5 | 100% | 11% |
| **m2** | 2.9 | | |
| **m3** | 3.6 | | |
| **m5** | 0.52 | | |
| **m6** | 2.80 | | |
| **m7** | 1.20 | | |
| **m8** | 1.30 | | |
| **m9** | 2.50 | | |
| **m10** | 1.30 | | |

%log: Percentage of mice that followed a logistic growth
%chaos: Percentage of mice following logistic growth that observed growth rate over 3.0

**Supplementary Table 8.** Lung cancer cell line models

Cell line: A549 (replicate 1)

| Control | Growth rate | %log | %chaos |
|---|---|---|---|
| **m1** | 0.7 | 88% | 0% |
| **m2** | 0.35 | | |
| **m3** | 0.9 | | |
| **m4** | 0.032 | | |
| **m5** | 0.66 | | |
| **m6** | 0.99 | | |
| **m7** | 0.74 | | |
| **m8** | n.a | | |

Cell line: A549 (replicate 2)

| Control | Growth rate | %log | %chaos |
|---|---|---|---|
| **m1** | 0.37 | 100% | 0% |
| **m2** | 0.6 | | |
| **m3** | 0.93 | | |
| **m4** | 0.62 | | |
| **m5** | 0.33 | | |
| **m6** | 0.25 | | |
| **m7** | 0.64 | | |
| **m8** | 0.49 | | |

Cell line: H520 (replicate 1)

| Control | Growth rate | %log | %chaos |
|---|---|---|---|
| **m1** | n.a | 60% | 0% |
| **m2** | 1.1 | | |
| **m3** | 2 | | |
| **m4** | n.a | | |
| **m5** | 1.6 | | |

Cell line: H520 (replicate 2)

| Control | Growth rate | %log | %chaos |
|---|---|---|---|
| **m1** | n.a | 40% | 0% |
| **m2** | 1.60 | | |
| **m3** | 1.70 | | |
| **m4** | n.a | | |
| **m5** | n.a | | |

Cell line: H441

| Control | Growth rate | %log | %chaos |
|---|---|---|---|
| **m1** | 0.65 | 75% | 0% |
| **m2** | 0.29 | | |
| **m3** | n.a | | |
| **m4** | 1.7 | | |
| **m5** | 2.1 | | |
| **m6** | 1.6 | | |
| **m7** | 0.33 | | |
| **m8** | n.a | | |

%log: Percentage of mice that followed a logistic growth
%chaos: Percentage of mice following logistic growth that observed growth rate over 3.0

**Supplementary Table 9.** Summary of growth rates across experiments. The growth rates are accumulated for all mice that adhered to logistic pattern. All growth rates more than 3 are color coded in orange to reflect possible mechanisms of chaos in those mice and all less than 3 are shown in green as these are unlikely to exhibit chaotic fluctuations. This heatmap provides an overview for possible inclination of cancer types under consideration to observe chaotic growth.

| Source | Cancer type | Growth rates | | | | | | | | |
|---|---|---|---|---|---|---|---|---|---|---|
| Hansson et.al. | Neuroblastoma | 4.1 | 4.4 | 4.9 | | | | | | |
| | Neuroblastoma | 2.9 | 6.9 | 7.1 | | | | | | |
| Radke et.al. | Neuroblastoma | 2.5 | 9.8 | 8.8 | | | | | | |
| Mohlin et.al. | Neuroblastoma | 4.8 | 6.2 | 5.8 | 1.7 | 11 | | | | |
| | Neuroblastoma | 1.4 | 3.7 | 4 | | | | | | |
| | Neuroblastoma | 5 | 1.5 | 5.6 | 3.7 | | | | | |
| | Neuroblastoma | 6.5 | 20 | 15 | | | | | | |
| | Neuroblastoma | 3.9 | 4.1 | | | | | | | |
| Manas et. al. | Neuroblastoma | 4.5 | 3.1 | 11 | | | | | | |
| | Neuroblastoma | 25 | 29 | 6 | | | | | | |
| | Neuroblastoma | 4.4 | 13 | 11 | 16 | 12 | | | | |
| | Neuroblastoma | 1.8 | 6.4 | | | | | | | |
| | Neuroblastoma | 9.4 | 9.8 | | | | | | | |
| | Neuroblastoma | 17 | 18 | 16 | 16 | | | | | |
| Murphy et.al. | Wilms tumor | 30 | 18 | 34 | 32 | 40 | 27 | | | |
| | Wilms tumor | 31 | 35 | 37 | 34 | 32 | 32 | 27 | 32 | |
| | Wilms tumor | 21 | 6.8 | 15 | 15 | 40 | 39 | 15 | | |
| | Wilms tumor | 5.8 | 20 | 12 | 4.3 | 7.3 | | | | |
| | Wilms tumor | 8.2 | 4.8 | 7.6 | | | | | | |
| | Wilms tumor | 4.4 | | | | | | | | |
| n.a. | Breast cancer | 0.9 | 0.23 | 0.98 | 0.63 | 0.97 | 0.99 | 0.98 | 0.87 | |
| | Breast cancer | 1.5 | 2.9 | 3.6 | 0.52 | 2.80 | 1.20 | 1.30 | 2.50 | 1.30 |
| | Lung cancer | 0.7 | 0.35 | 0.9 | 0.032 | 0.66 | 0.99 | 0.74 | | |
| | Lung cancer | 0.37 | 0.6 | 0.93 | 0.62 | 0.33 | 0.25 | 0.64 | 0.49 | |
| | Lung cancer | 1.1 | 2 | 1.6 | | | | | | |
| | Lung cancer | 1.60 | 1.70 | | | | | | | |
| | Lung cancer | 0.65 | 0.29 | 1.7 | 2.1 | 1.6 | 0.33 | | | |

| scale |
|---|
| <1 |
| 3 |
| >4 |

### Supplementary figure legends

#### Supplementary figure 1-4.

Prediction of growth trend (red if logistic, green if not) are shown for each growth curve, experiment wise. For neuroblastoma, **a-e** are from Mohlin et. al., **f-g** are from Hansson et. al., **h** is from Radke et. al. and, **i-n** are from Manas et. al. The Wilms tumor, breast and lung cancer samples are taken from sources as discussed in supplementary table 1. All samples are ordered as they appear in supplementary tables 2-8.

#### Supplementary figures 5-30.

Good ness of logistic fit is shown with fitted curve and overlayed tumor volume measures. Each plot refers to a single control mouse belonging to a set of experiments whose growth adhered to a logistic growth. For example, mice from replicate 1 of Hansson et. al. are shown in supplementary figure 5. Supplementary table 2 for the replicate 1 shows three mice (A1, A2, A3) followed a logistic growth. In supplementary figure 5, goodness of fit are shown for these three mice. All plots accompany the mouse identifier that correspond to supplementary tables 2-8 for respective experiment.

Supplementary figure 1. Neuroblastoma

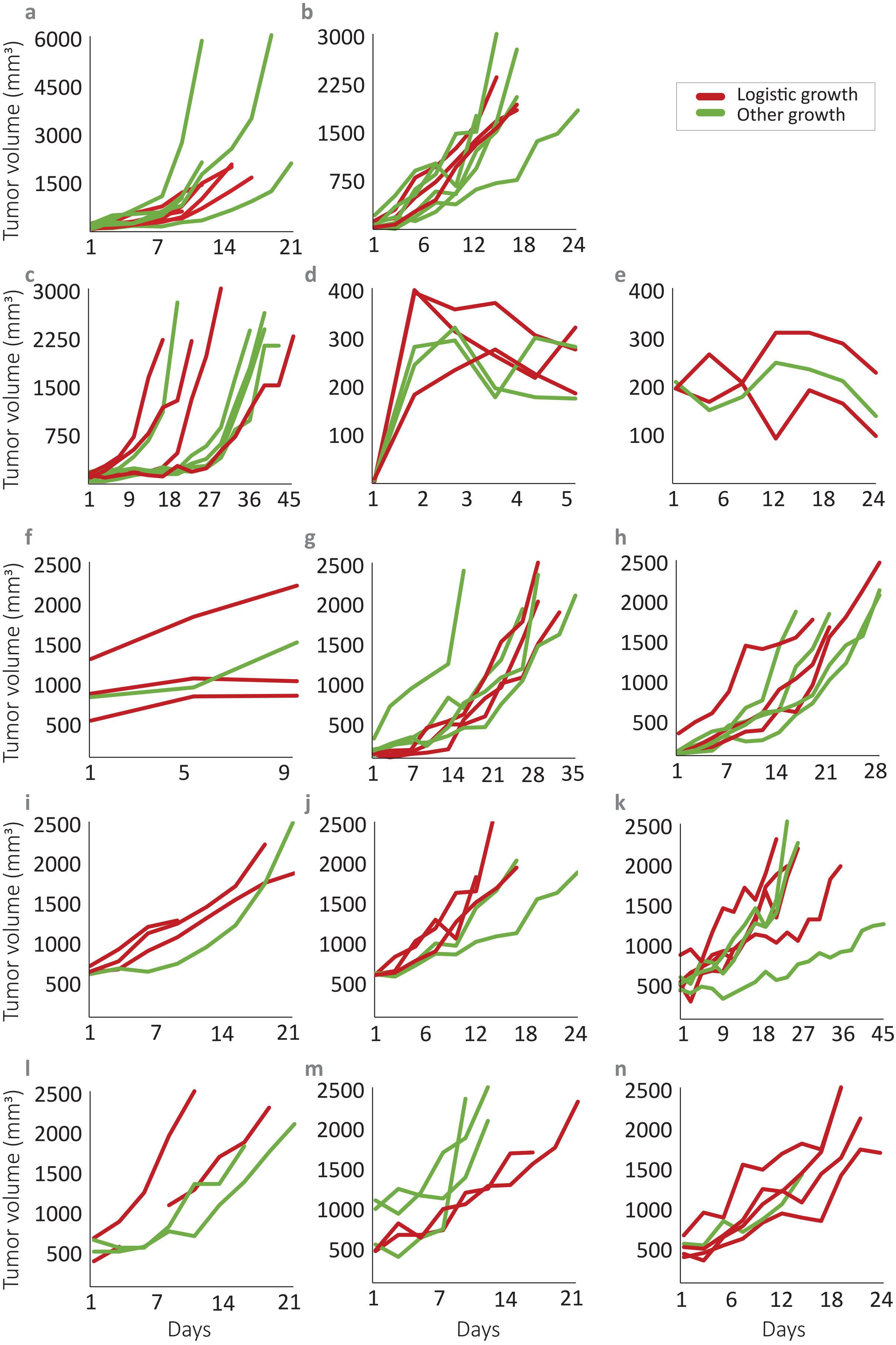

Supplementary figure 2. Wilms tumor

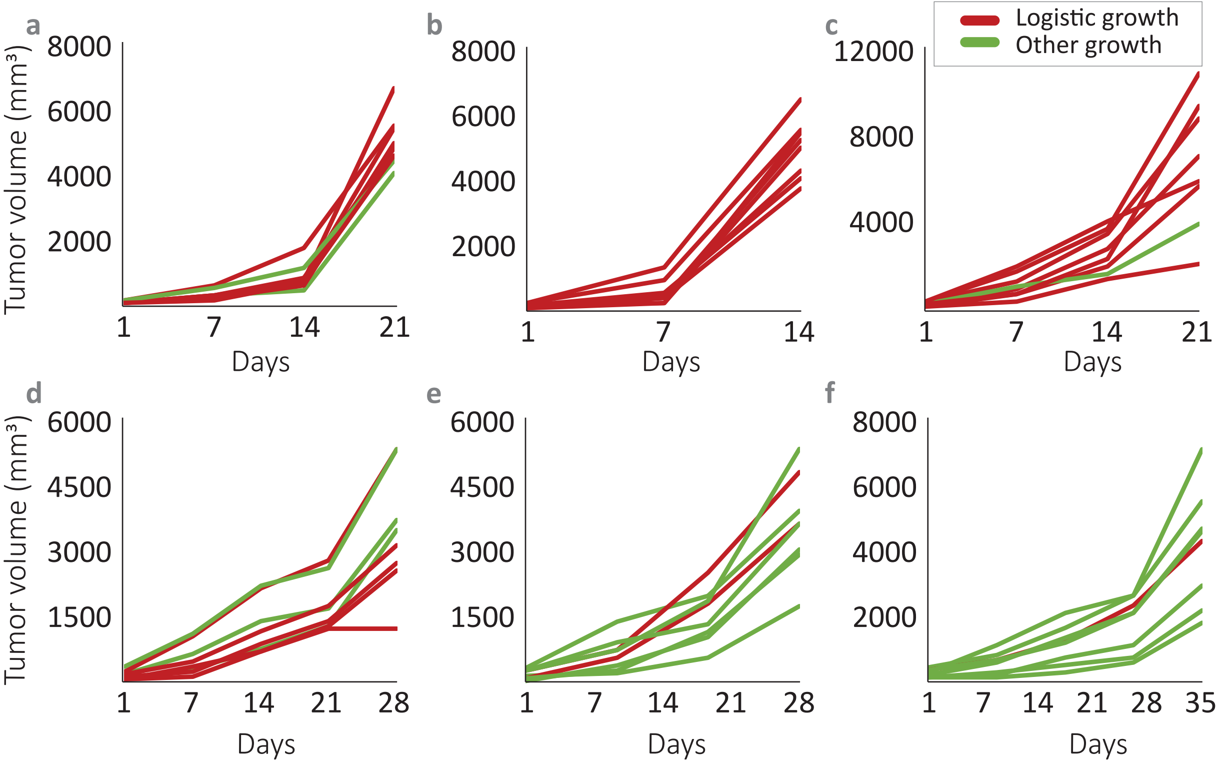

**Supplementary figure 3.** **Breast cancer**

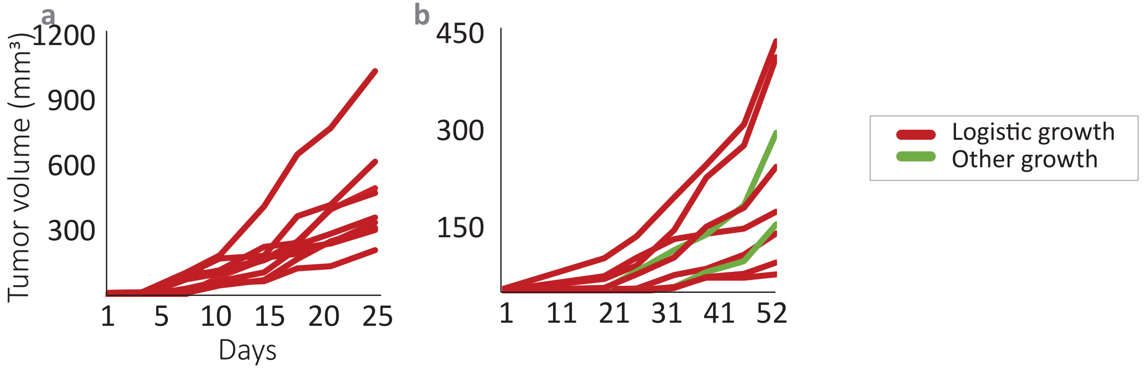


**Supplementary figure 4.** **Lung cancer**

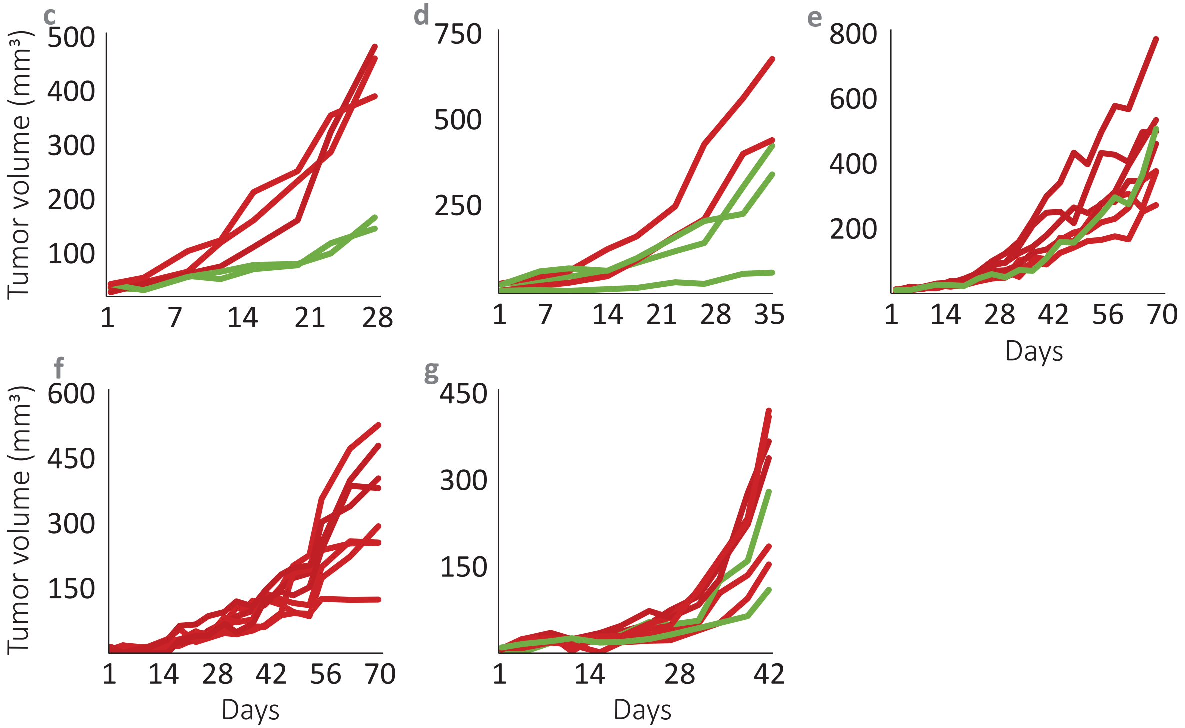

**Supplementary figure 5.** Neuroblastoma PDX (Hansson et. al., replicate 1)

mouse A1

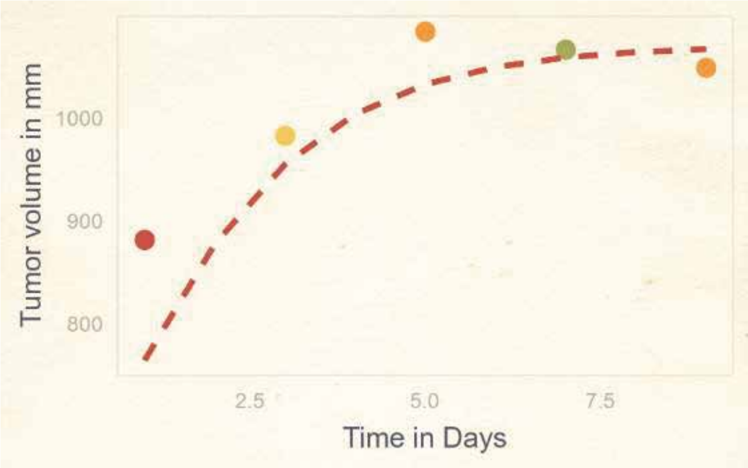


mouse A2

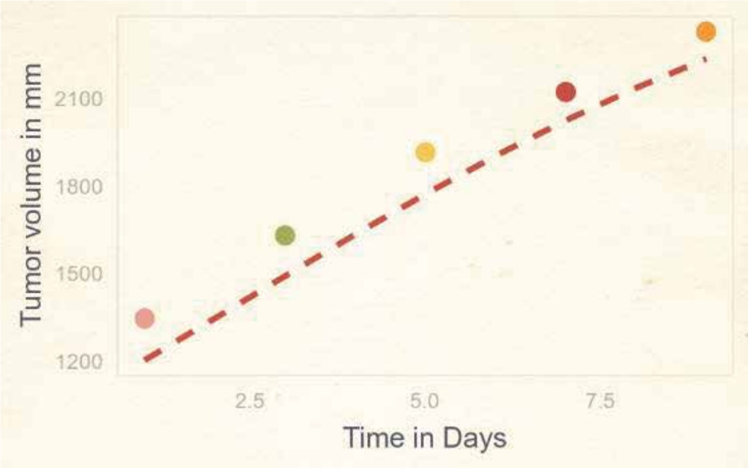


mouse A3

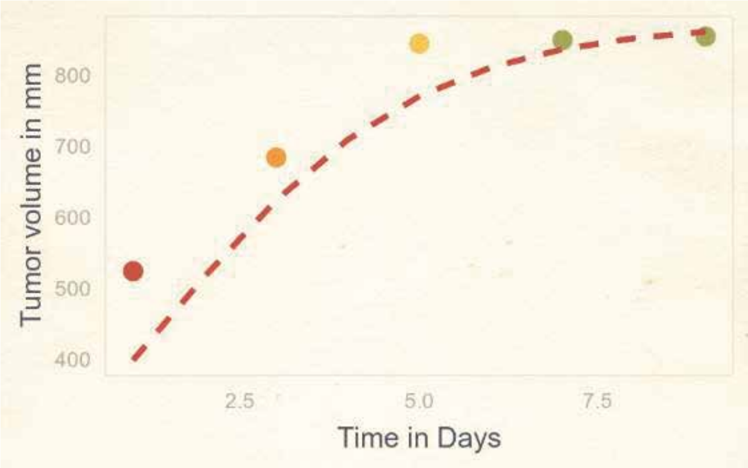


**Supplementary figure 6.** Neuroblastoma PDX (Hansson et. al., replicate 2)

mouse A1

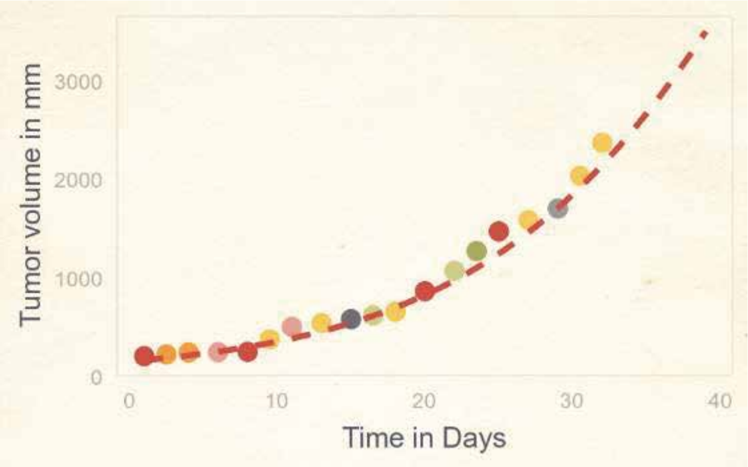


mouse A4

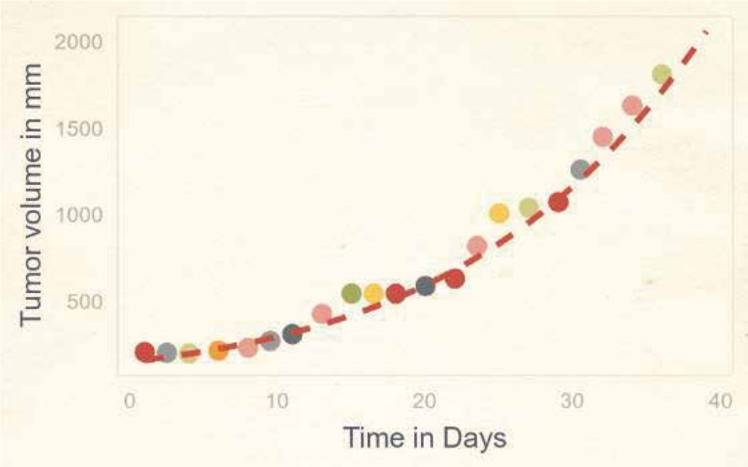


mouse B1

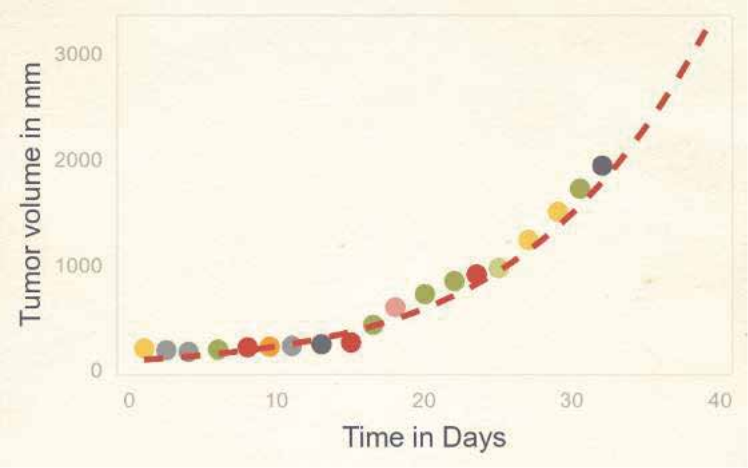

**Supplementary figure 7.** Neuroblastoma PDX (Radke et. al.)

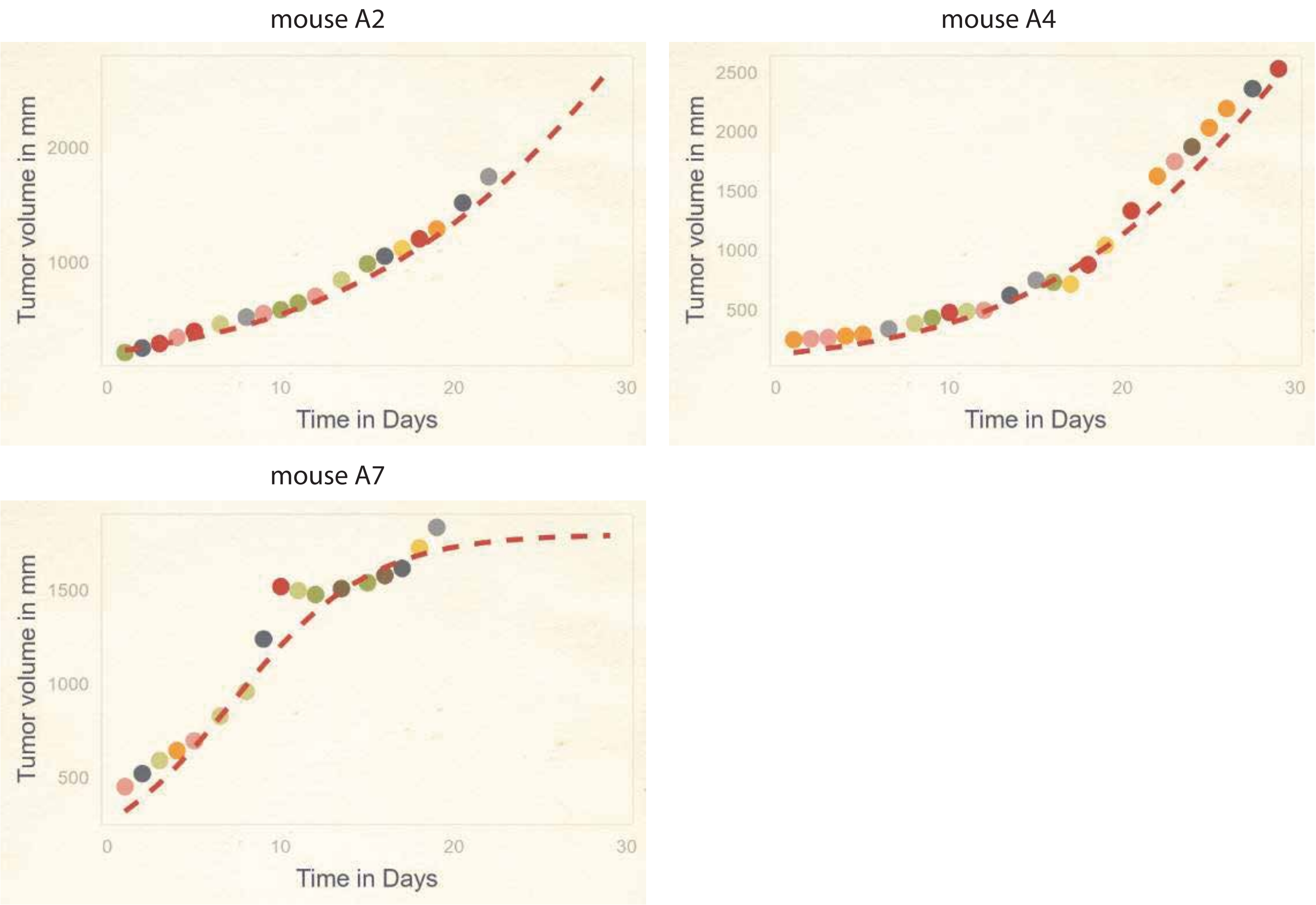

**Supplementary figure 8.** Neuroblastoma PDX (Mohlin et. al., replicate 1)

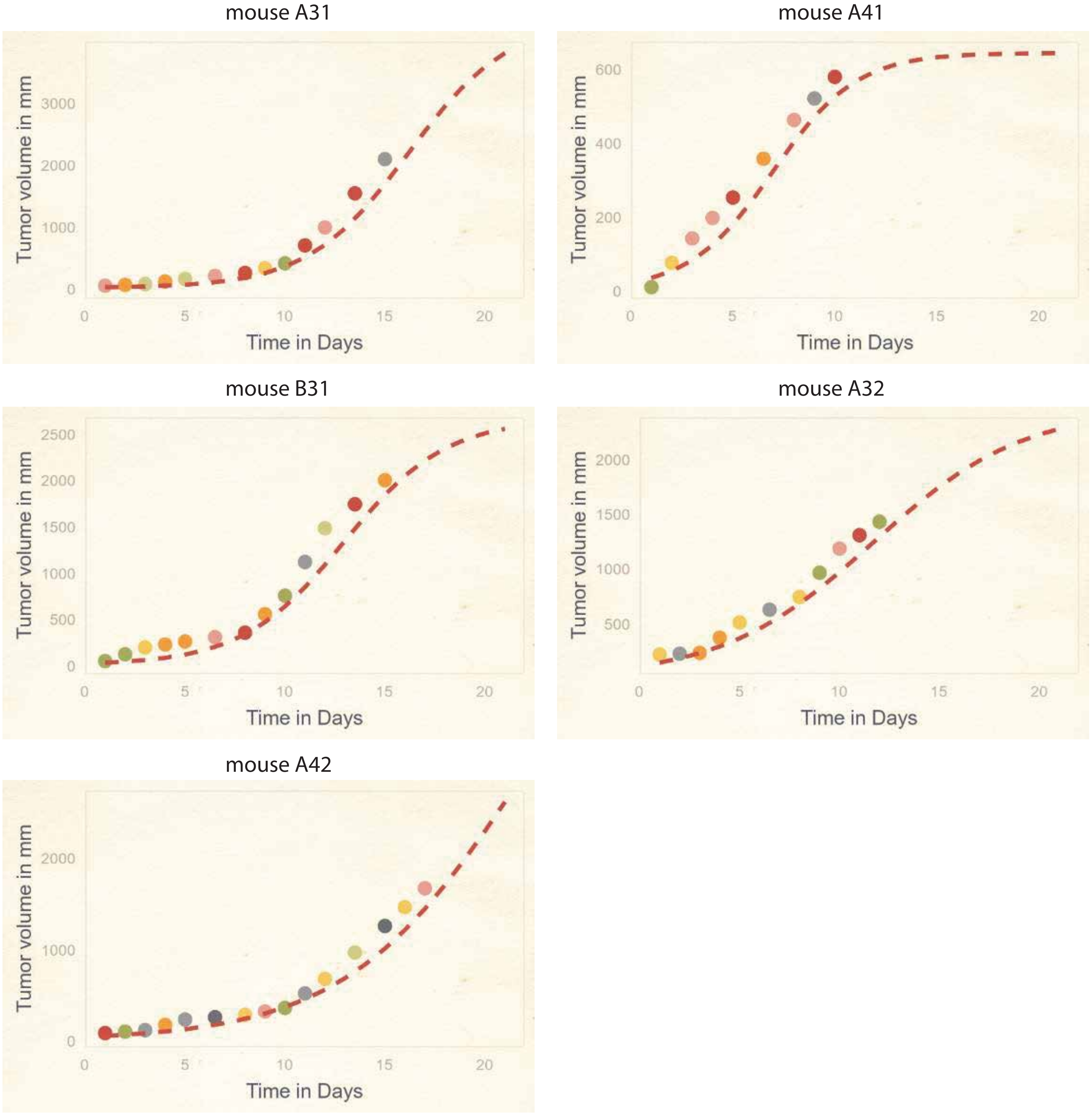


**Supplementary figure 9.** Neuroblastoma PDX (Mohlin et. al., replicate 2)

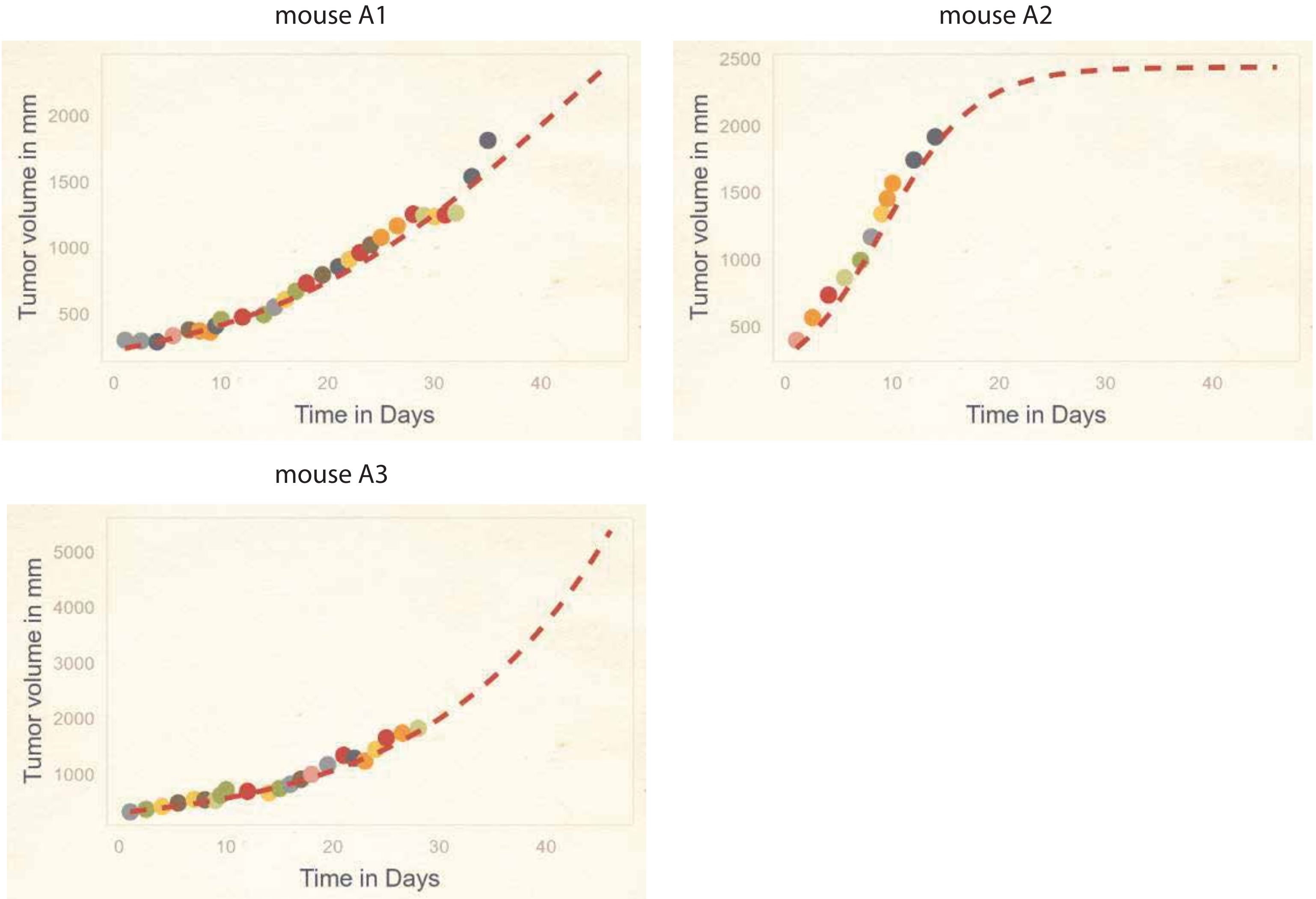

**Supplementary figure 10.** Neuroblastoma PDX (Mohlin et. al., replicate 3)

mouse 2

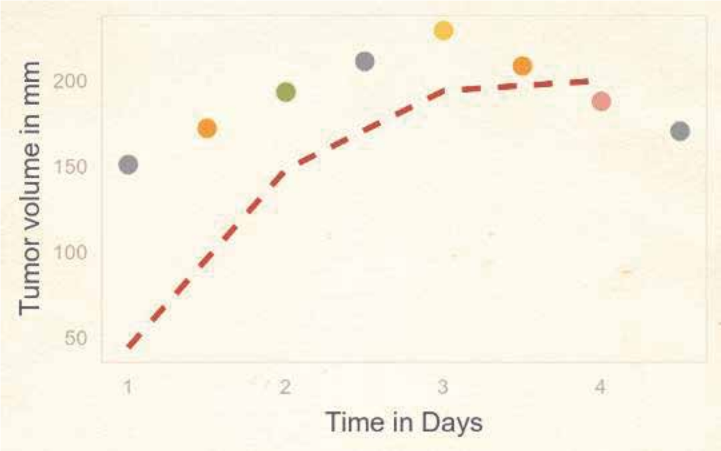


mouse 3

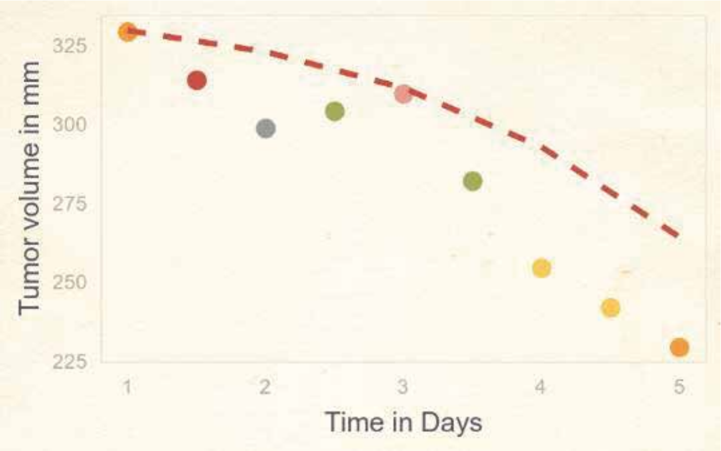


mouse 5

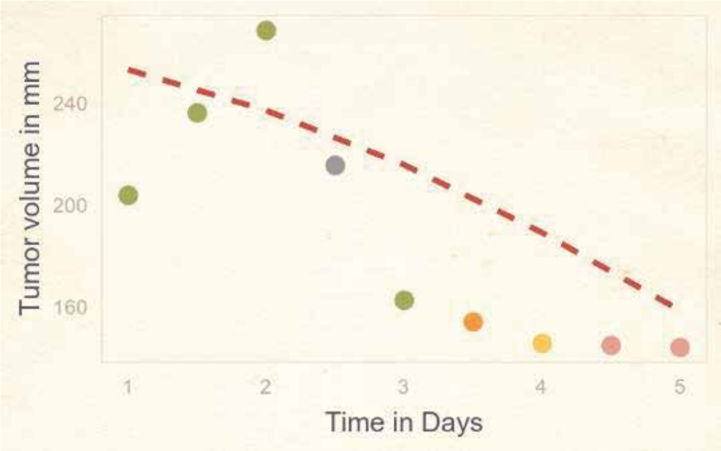


**Supplementary figure 11.** Neuroblastoma PDX (Mohlin et. al., replicate 4)

mouse 1

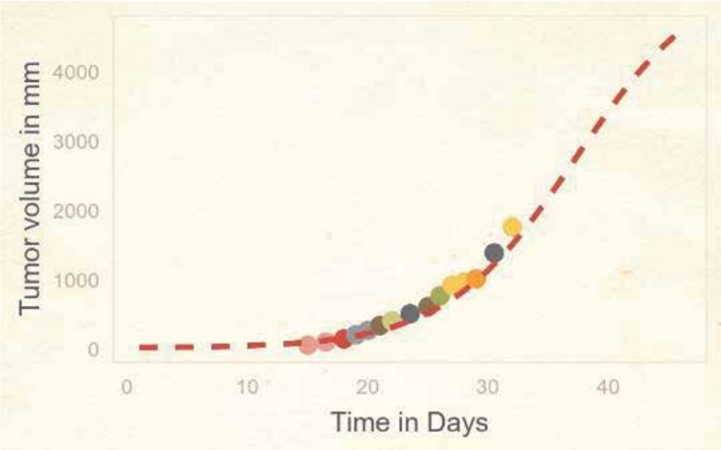


mouse 7

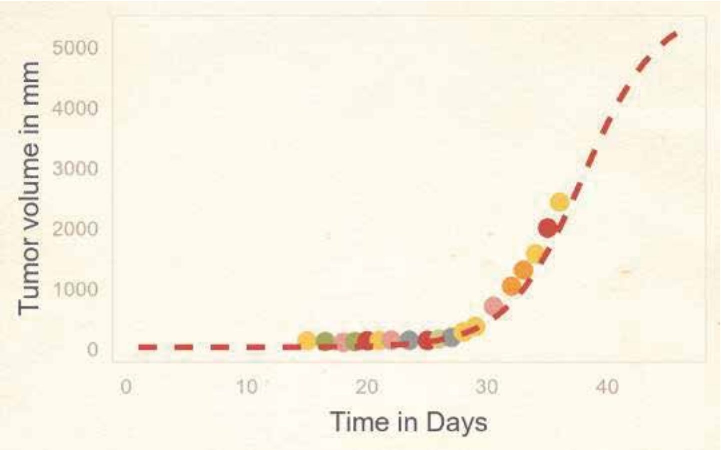


mouse 8

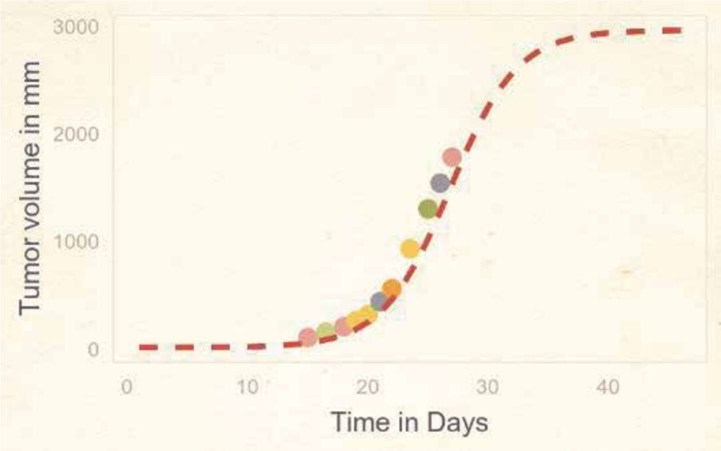


mouse 9

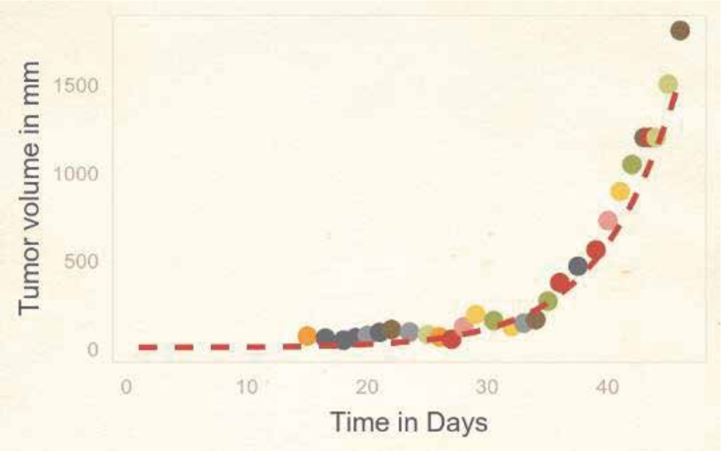


**Supplementary figure 12.** Neuroblastoma tumor initiating cells (Mohlin et. al.)

mouse 1

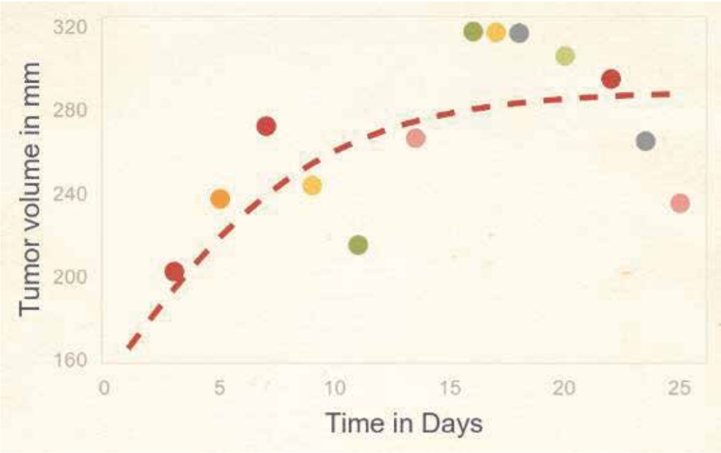


mouse 4

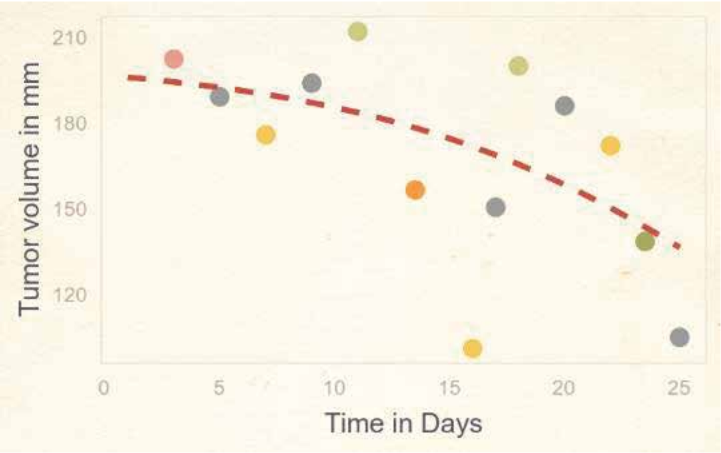

**Supplementary figure 13.** Neuroblastoma PDX (Manas et. al., *PDX1*, replicate 1)

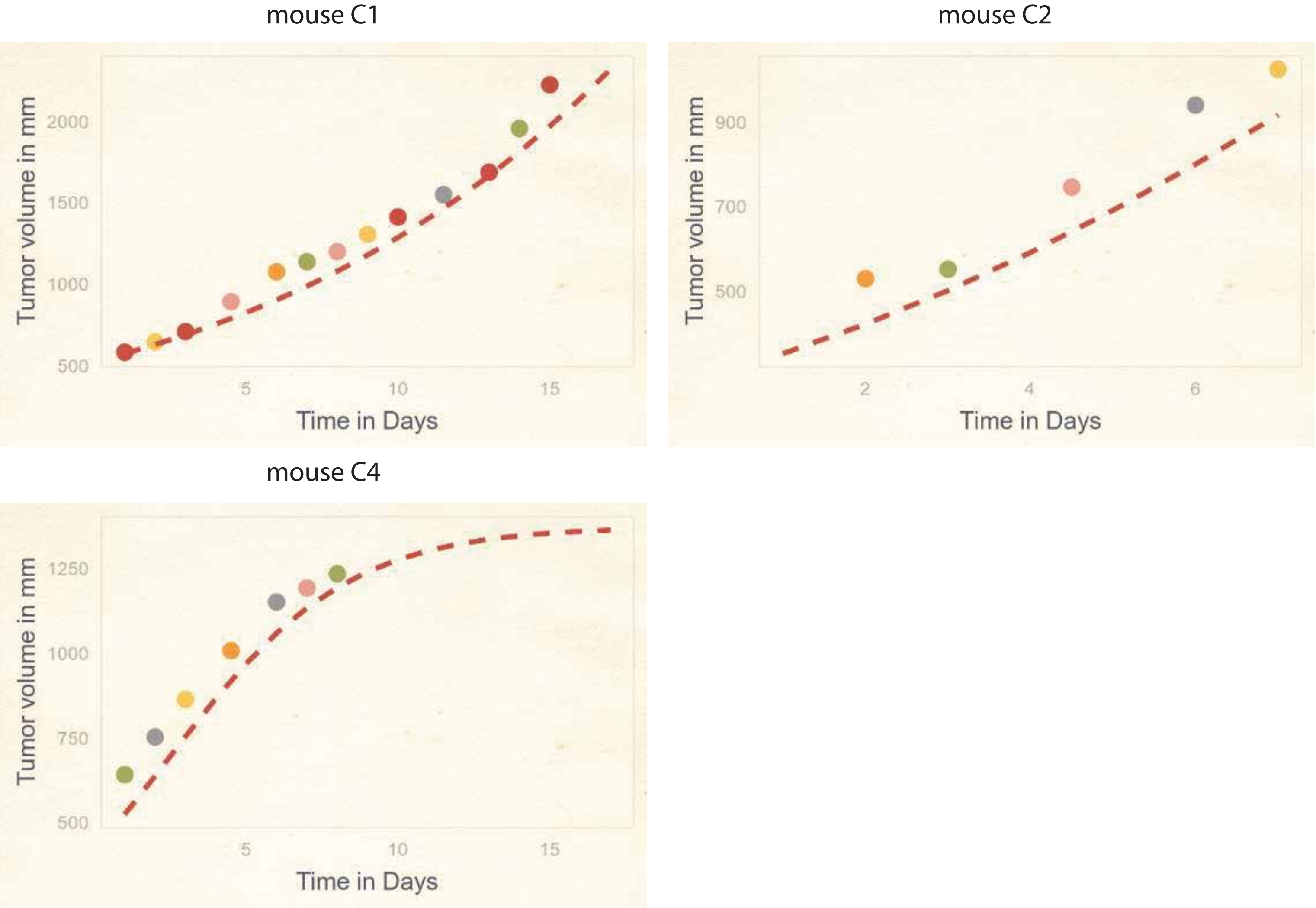


**Supplementary figure 14.** Neuroblastoma PDX (Manas et. al., *PDX1*, replicate 2)

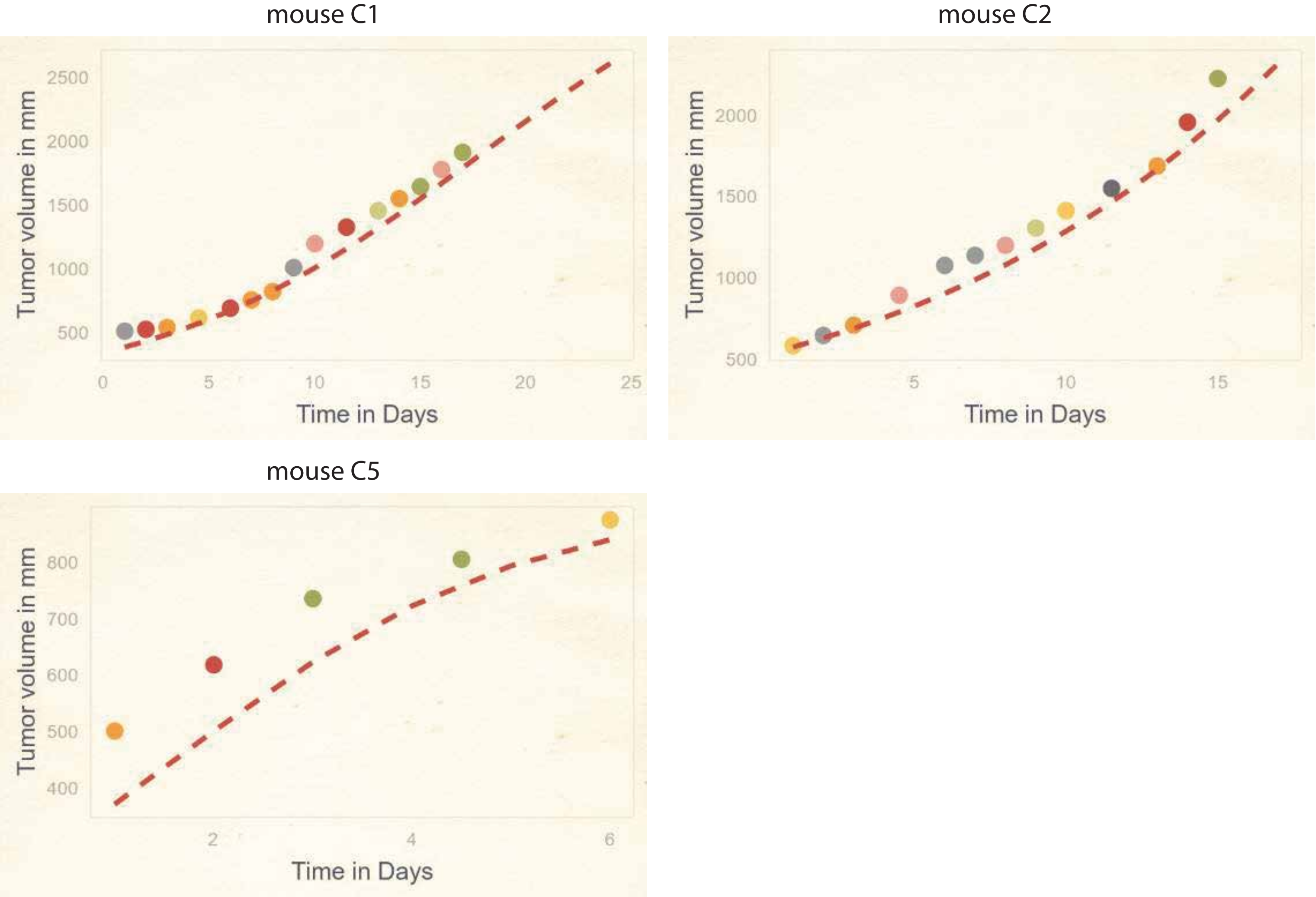

**Supplementary figure 15.** Neuroblastoma PDX (Manas et. al., *PDX2*)

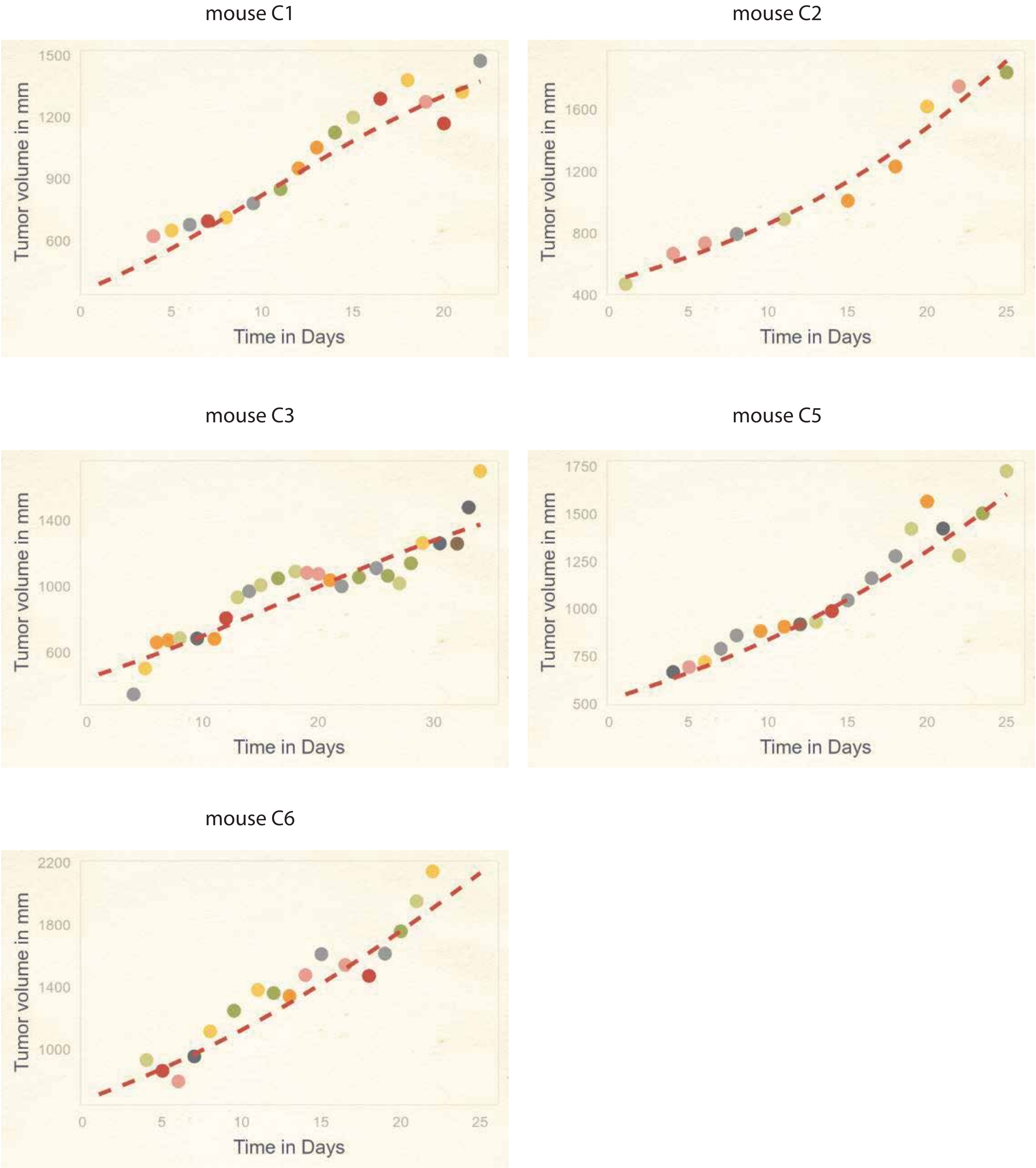

**Supplementary figure 16.** Neuroblastoma PDX (Manas et. al., *PDX3*, replicate 1)

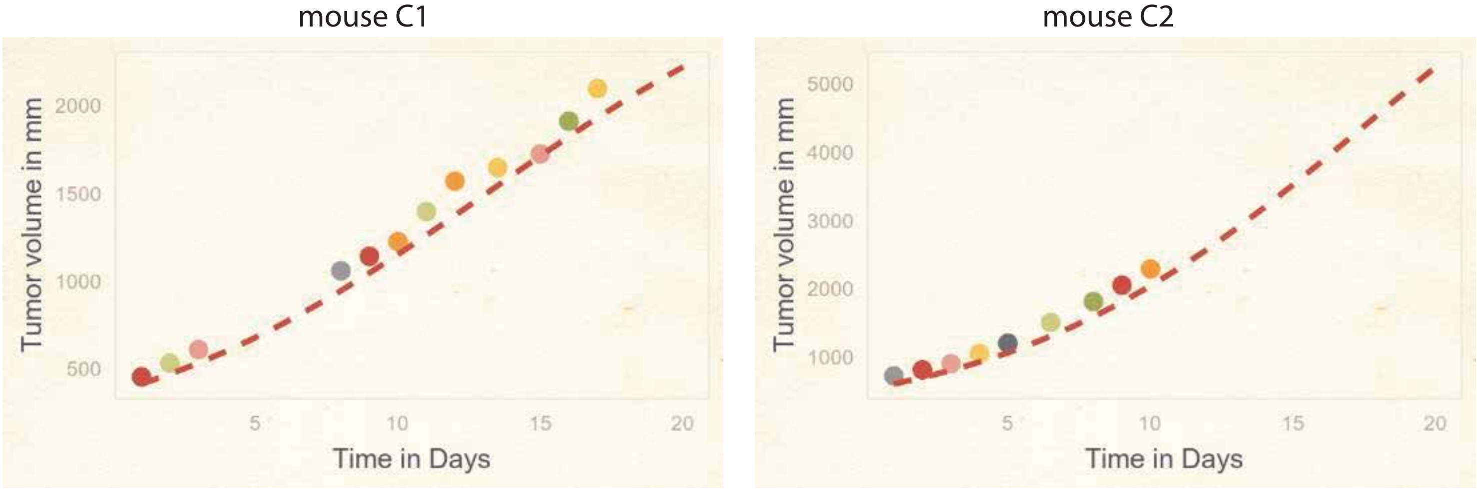


**Supplementary figure 17.** Neuroblastoma PDX (Manas et. al., *PDX3*, replicate 2)

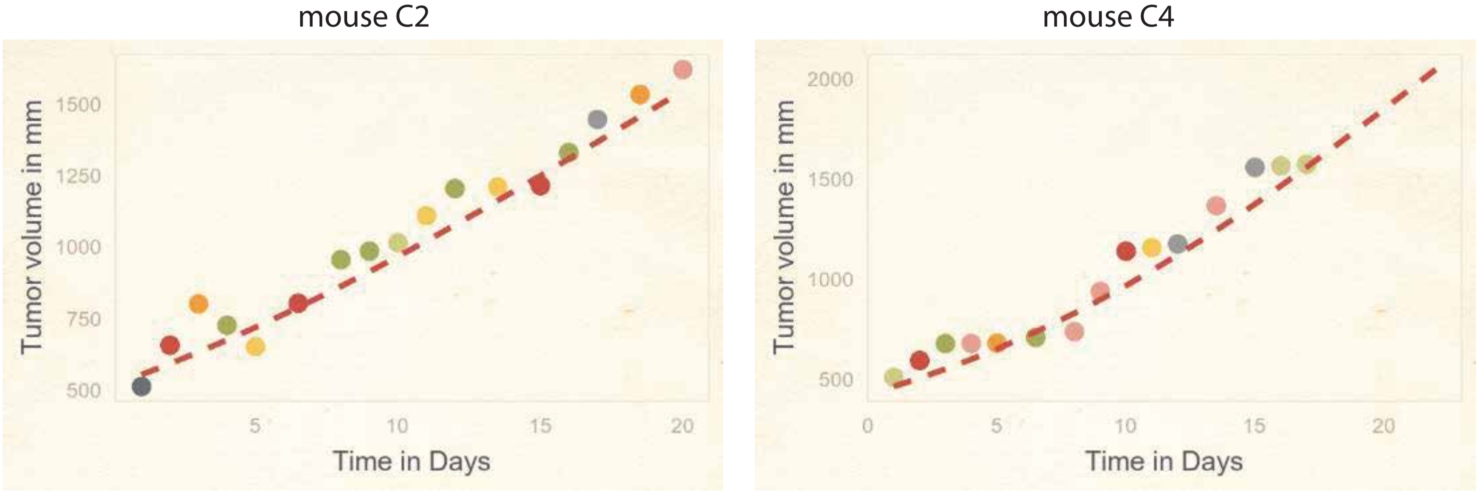


**Supplementary figure 18.** Neuroblastoma PDX (Manas et. al., *PDX3*, replicate 3)

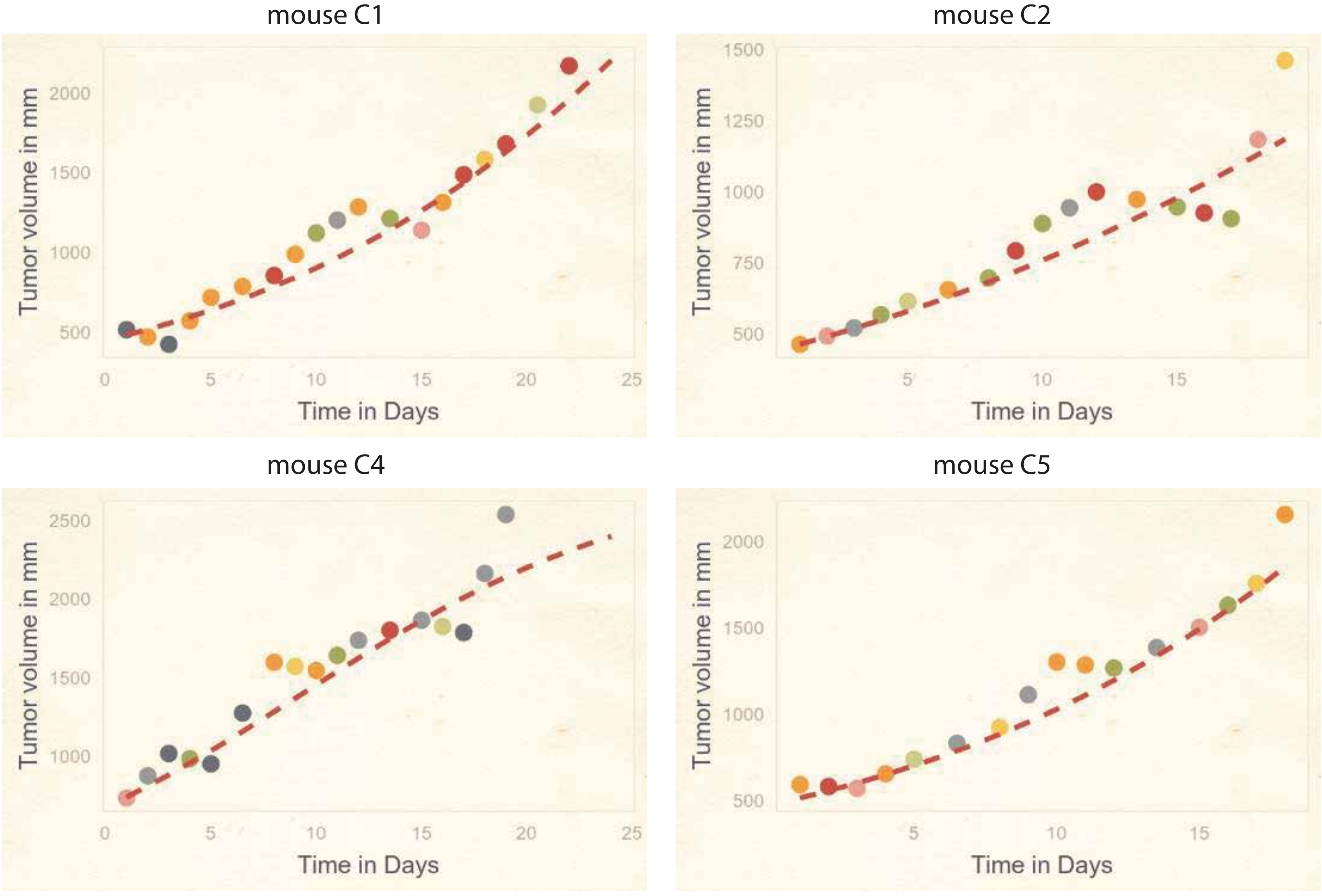

**Supplementary figure 19.** Wilms tumor PDX KT47

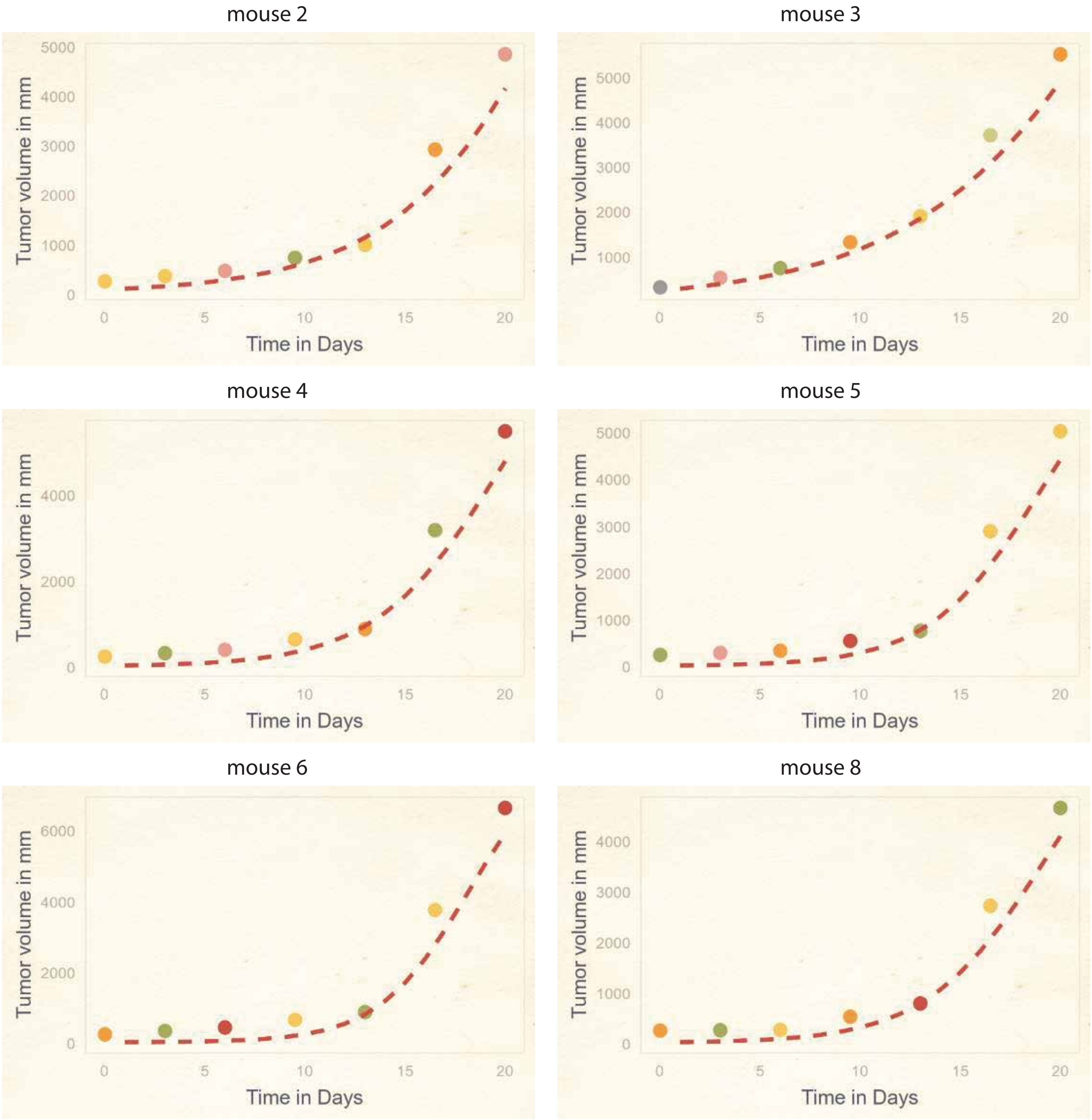

**Supplementary figure 20.** Wilms tumor PDX KT53

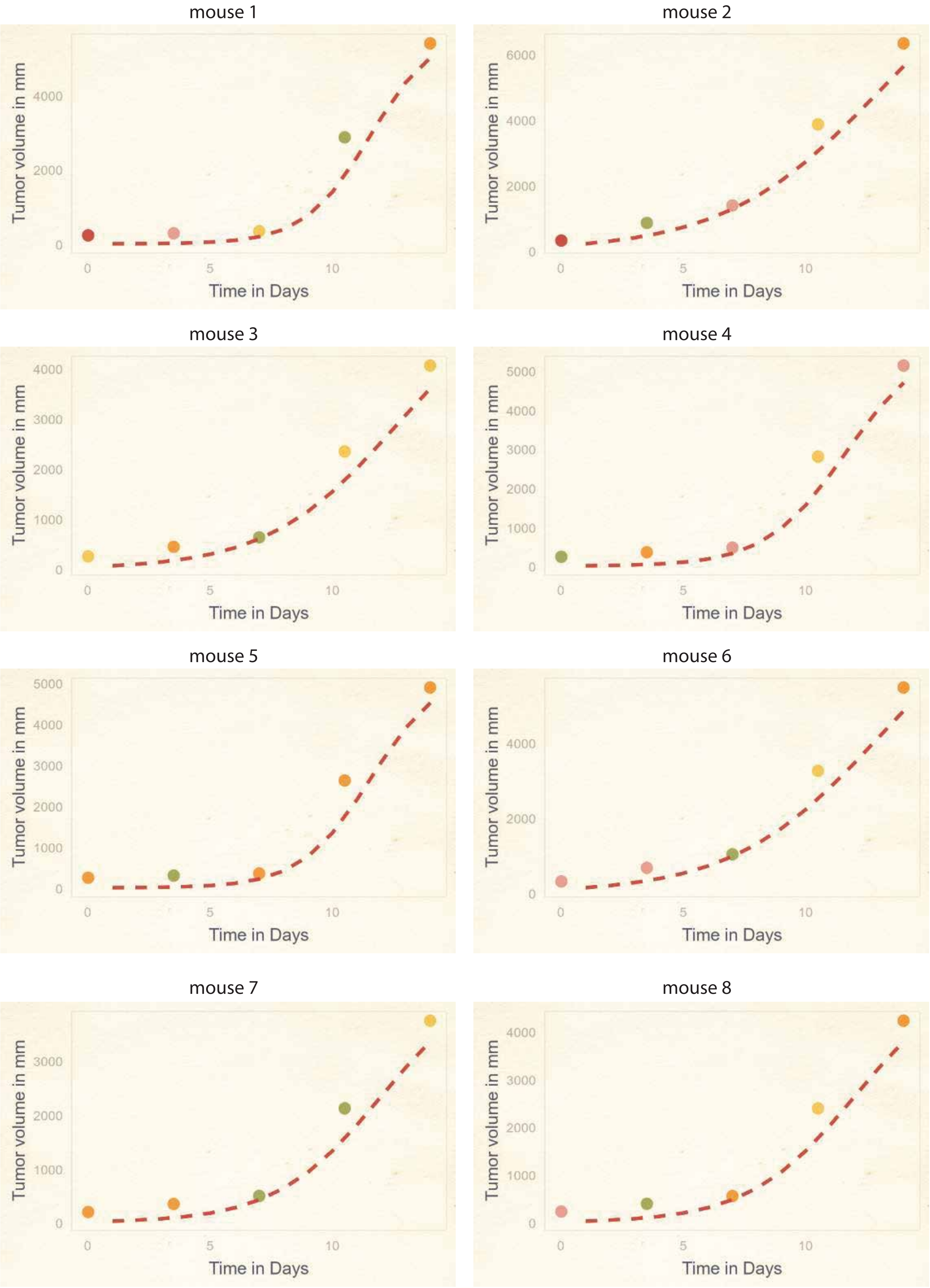

**Supplementary figure 21.** Wilms tumor PDX KT51

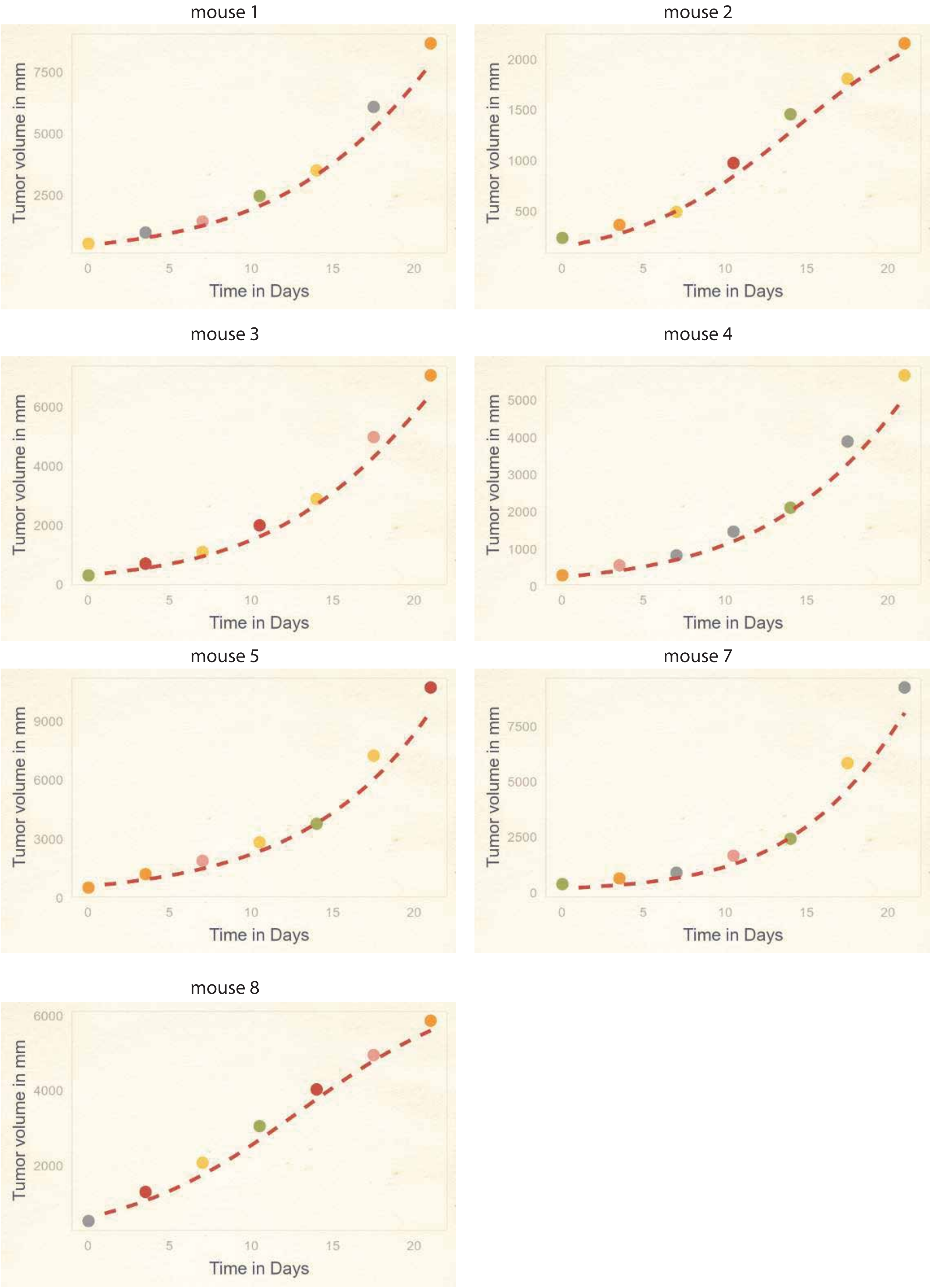

**Supplementary figure 22.** Wilms tumor PDX KT75

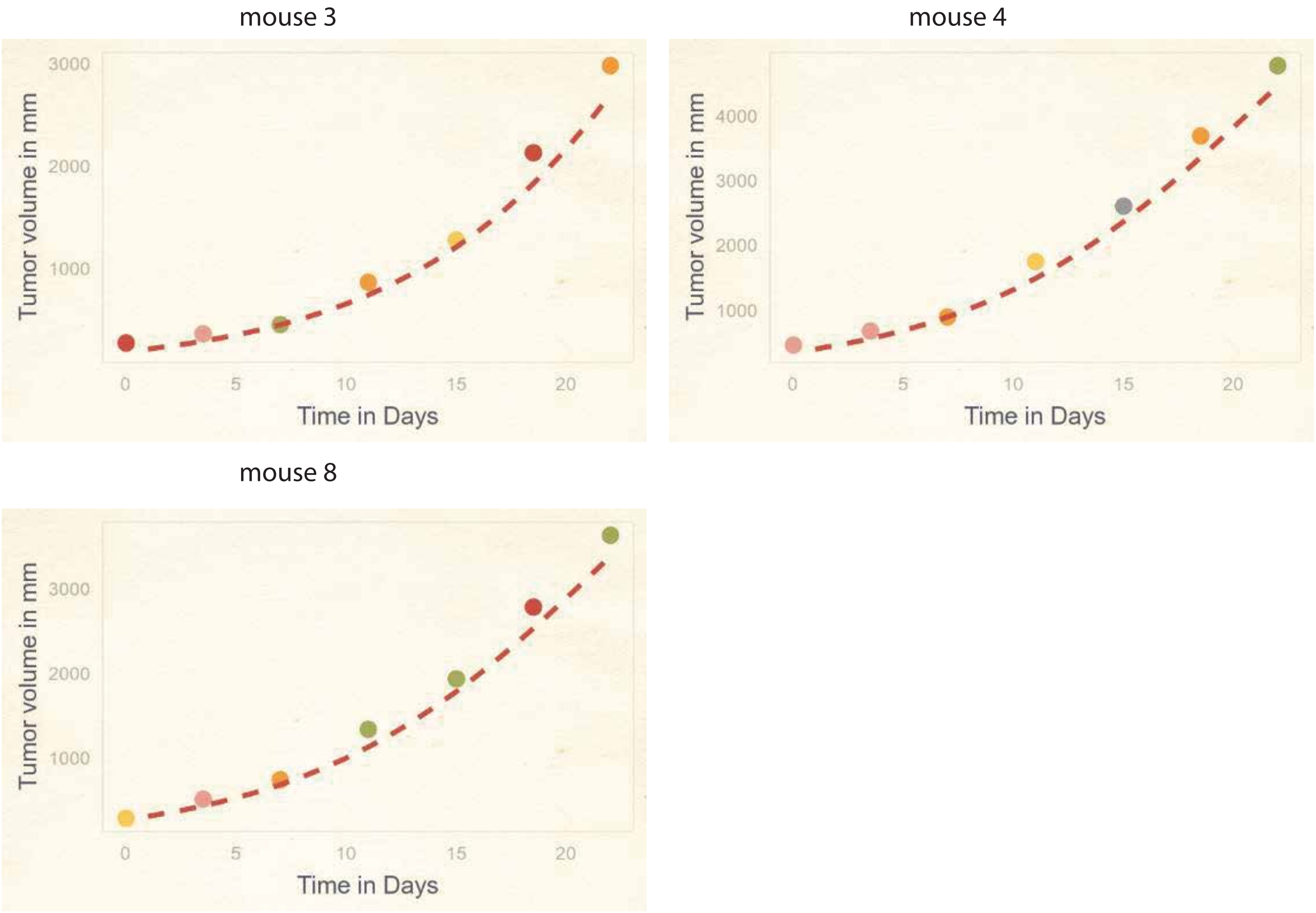


**Supplementary figure 23.** Wilms tumor PDX KT43

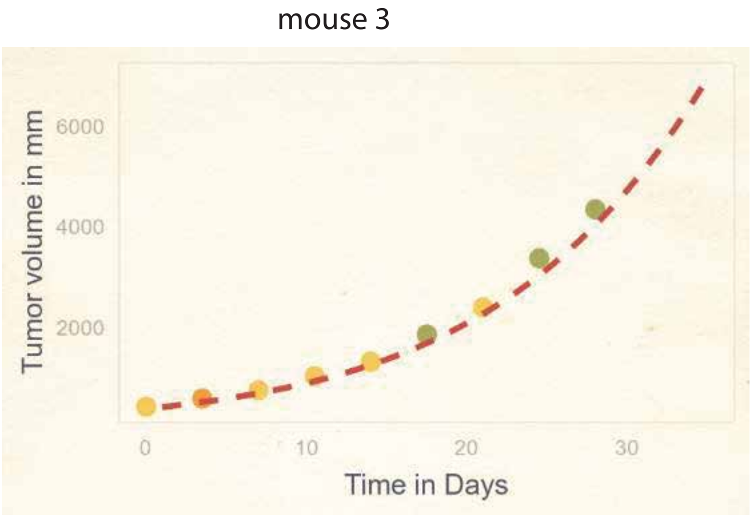

**Supplementary figure 24.** A549 lung cancer cell line (replicate 1)

mouse 1

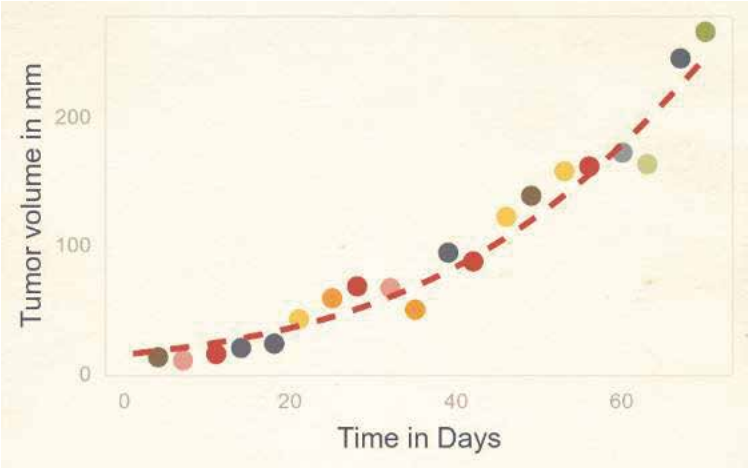


mouse 2

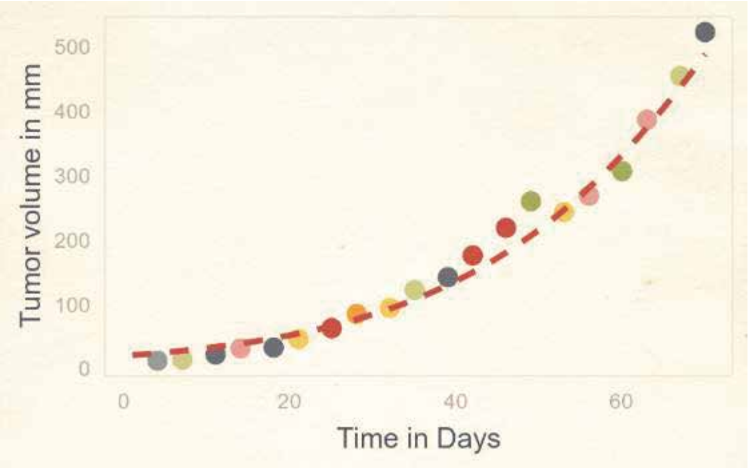


mouse 4

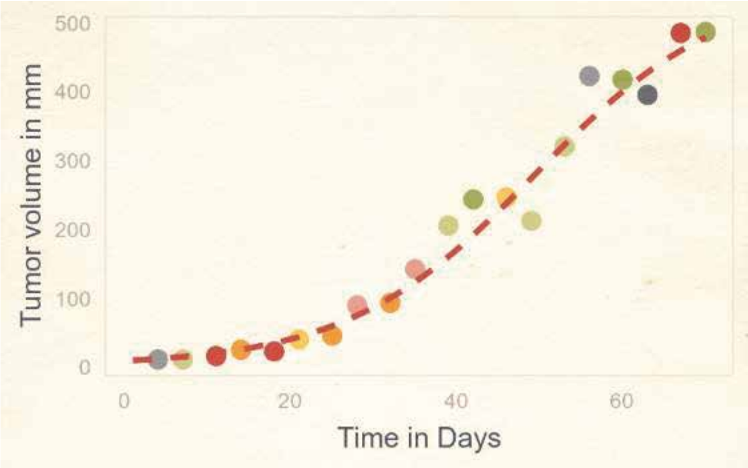


mouse 5

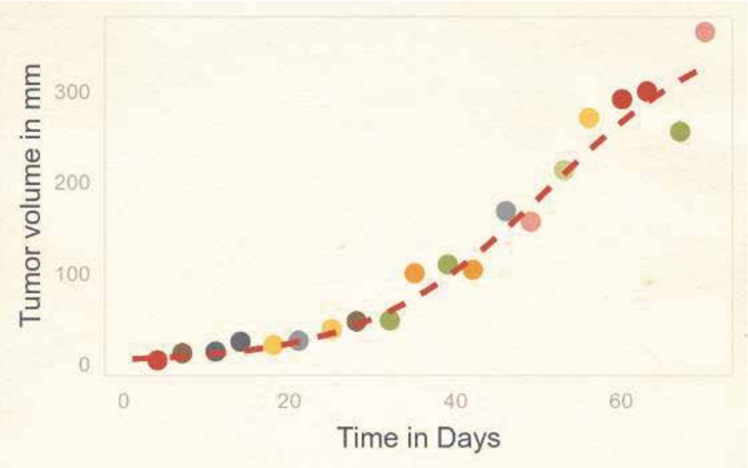


mouse 6

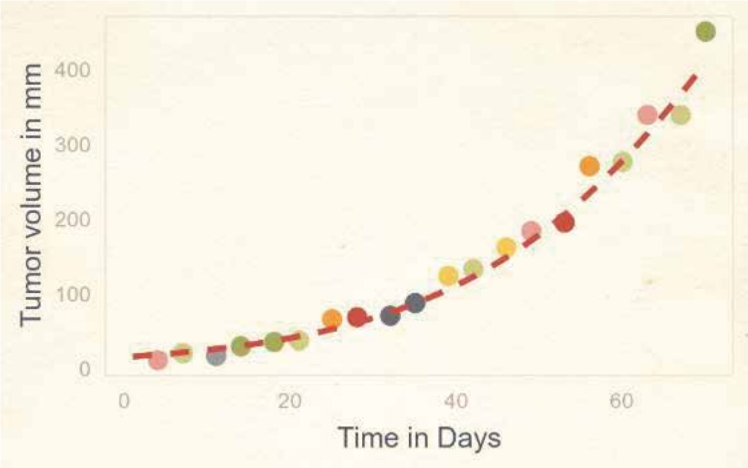


mouse 7

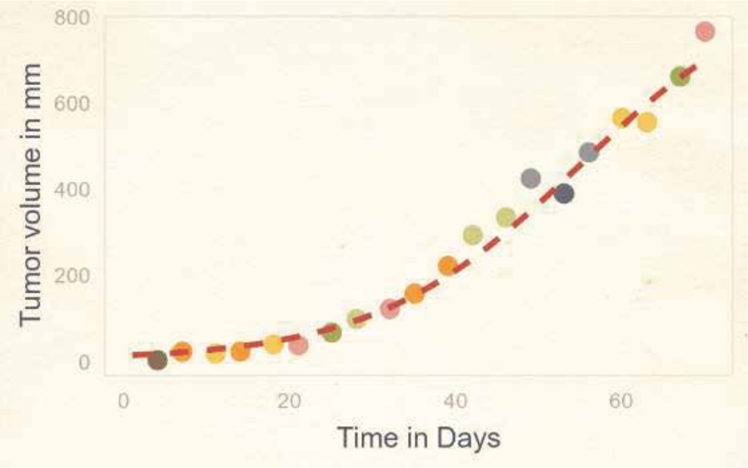


mouse 7

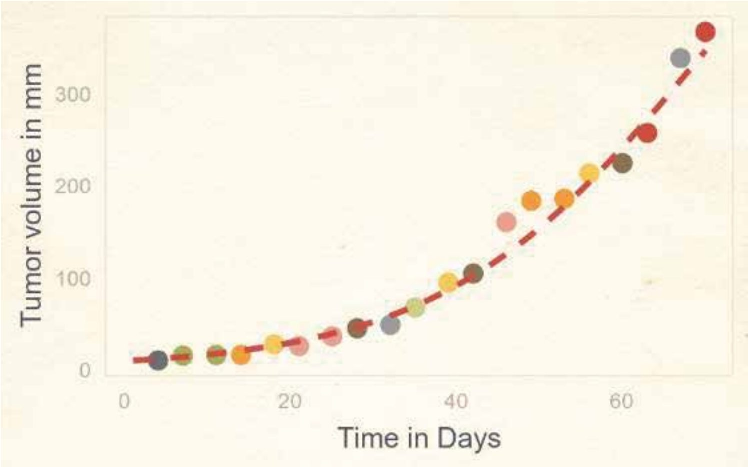

**Supplementary figure 25.** A549 lung cancer cell line (replicate 2)

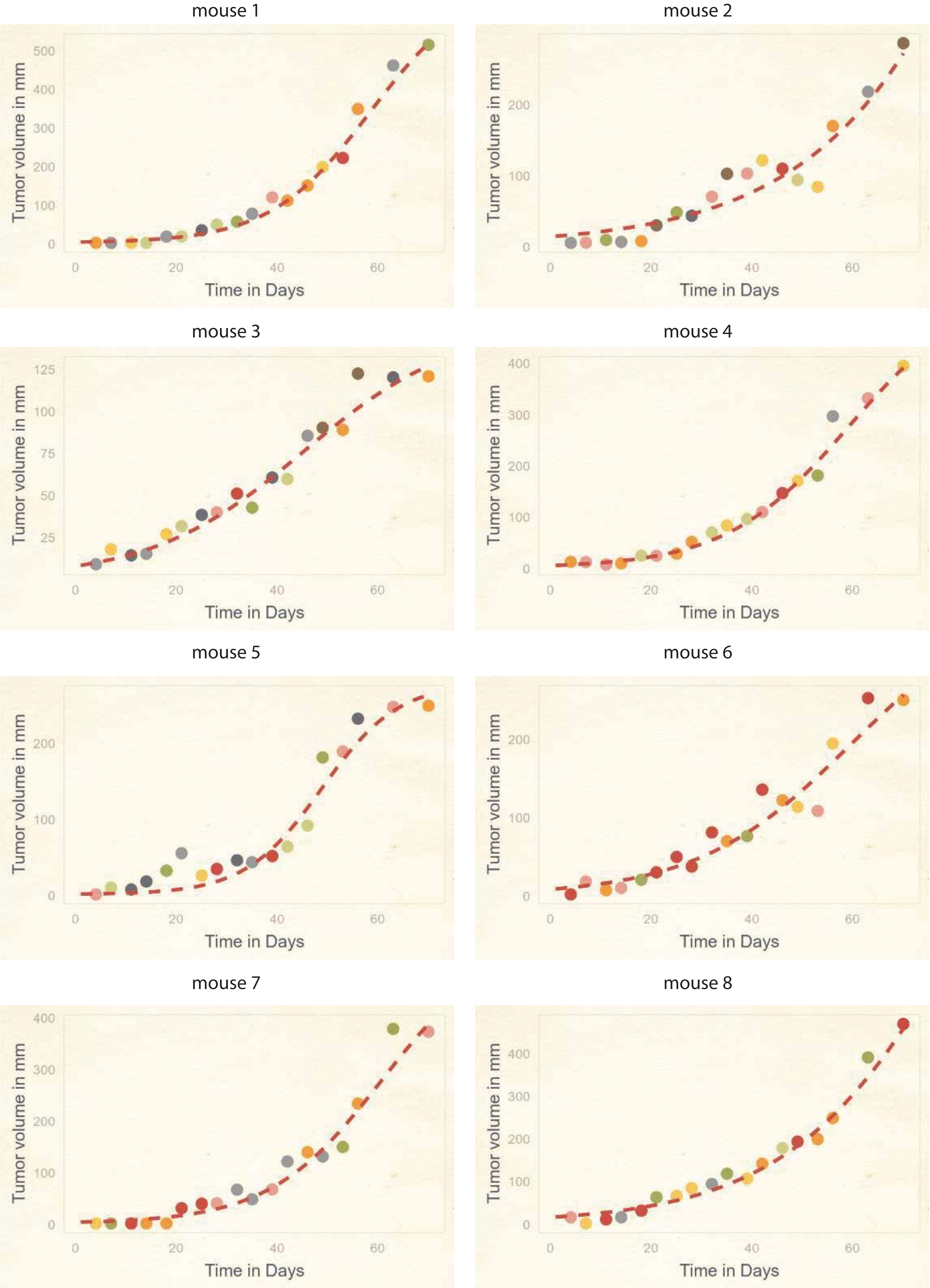

**Supplementary figure 26.** H441 lung cancer cell line

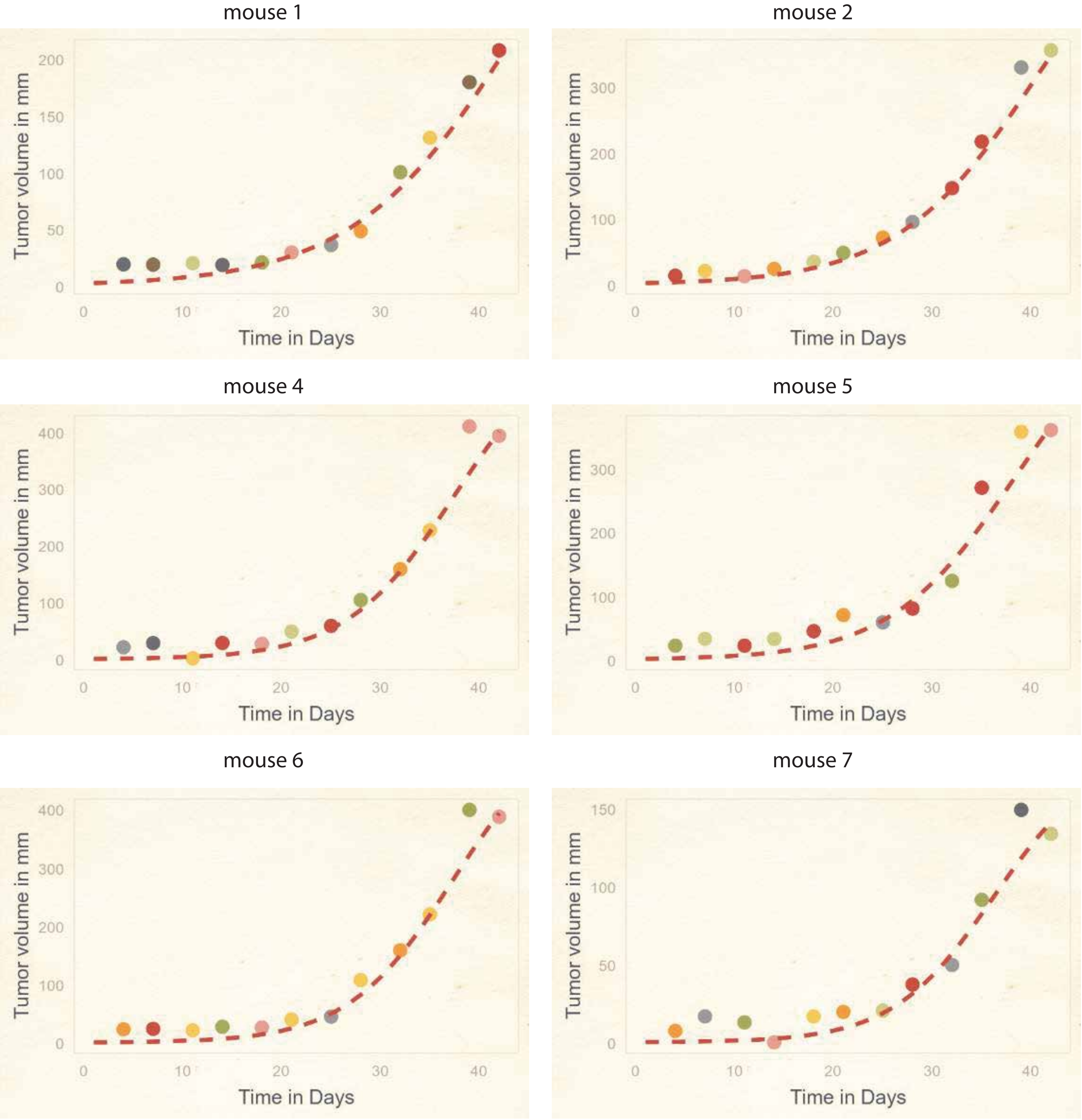

**Supplementary figure 27.** H520 lung cancer cell line (replicate 1)

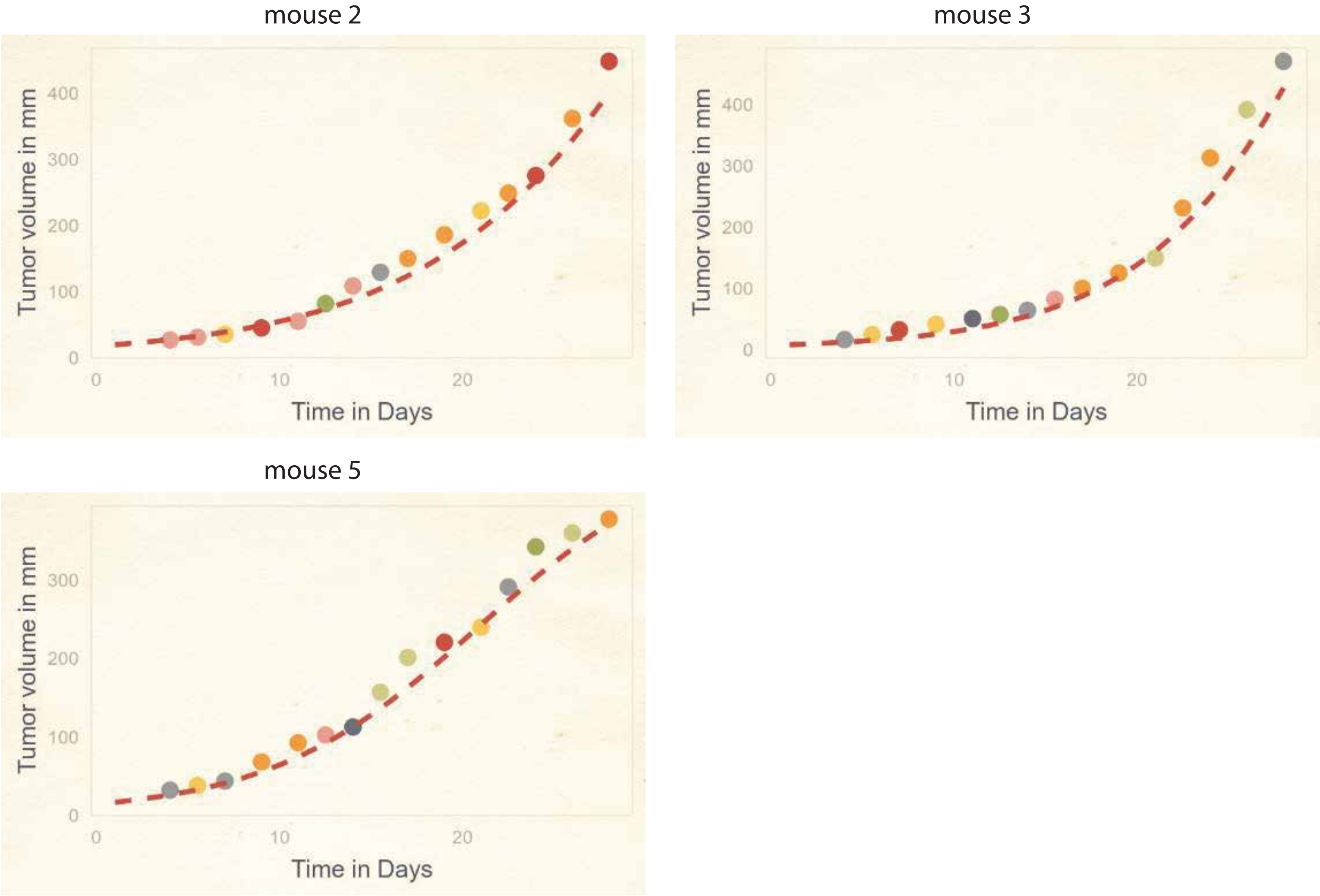


**Supplementary figure 28.** H520 lung cancer cell line (replicate 2)

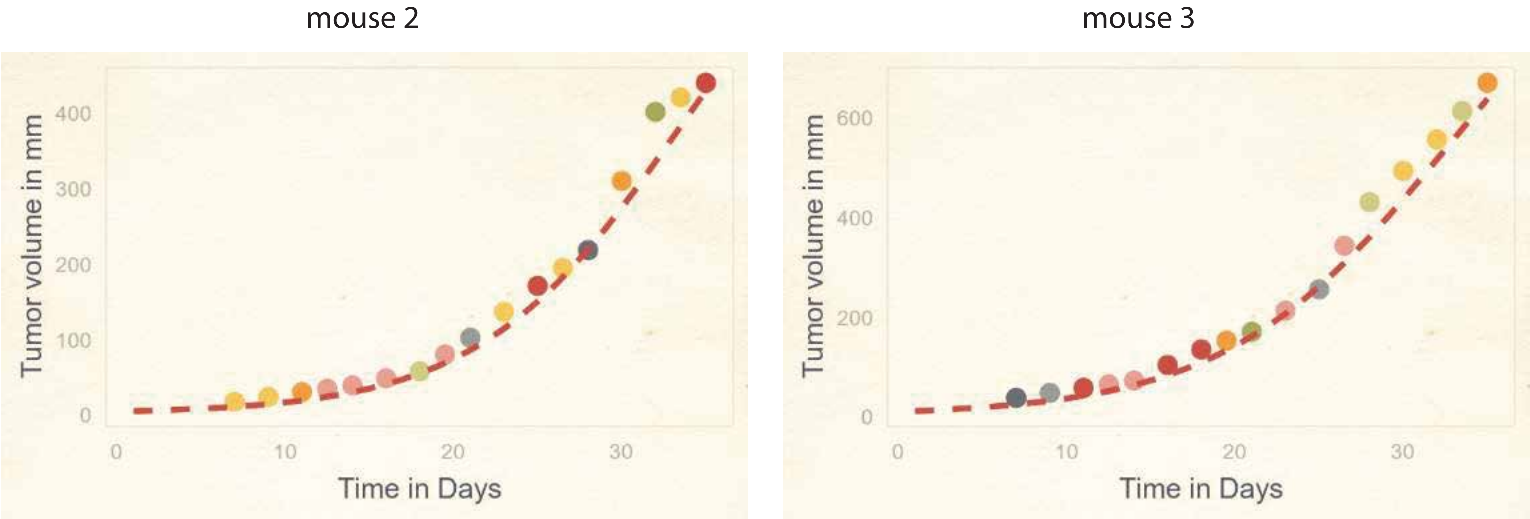

**Supplementary figure 29.** MCF7 breast cancer cell line

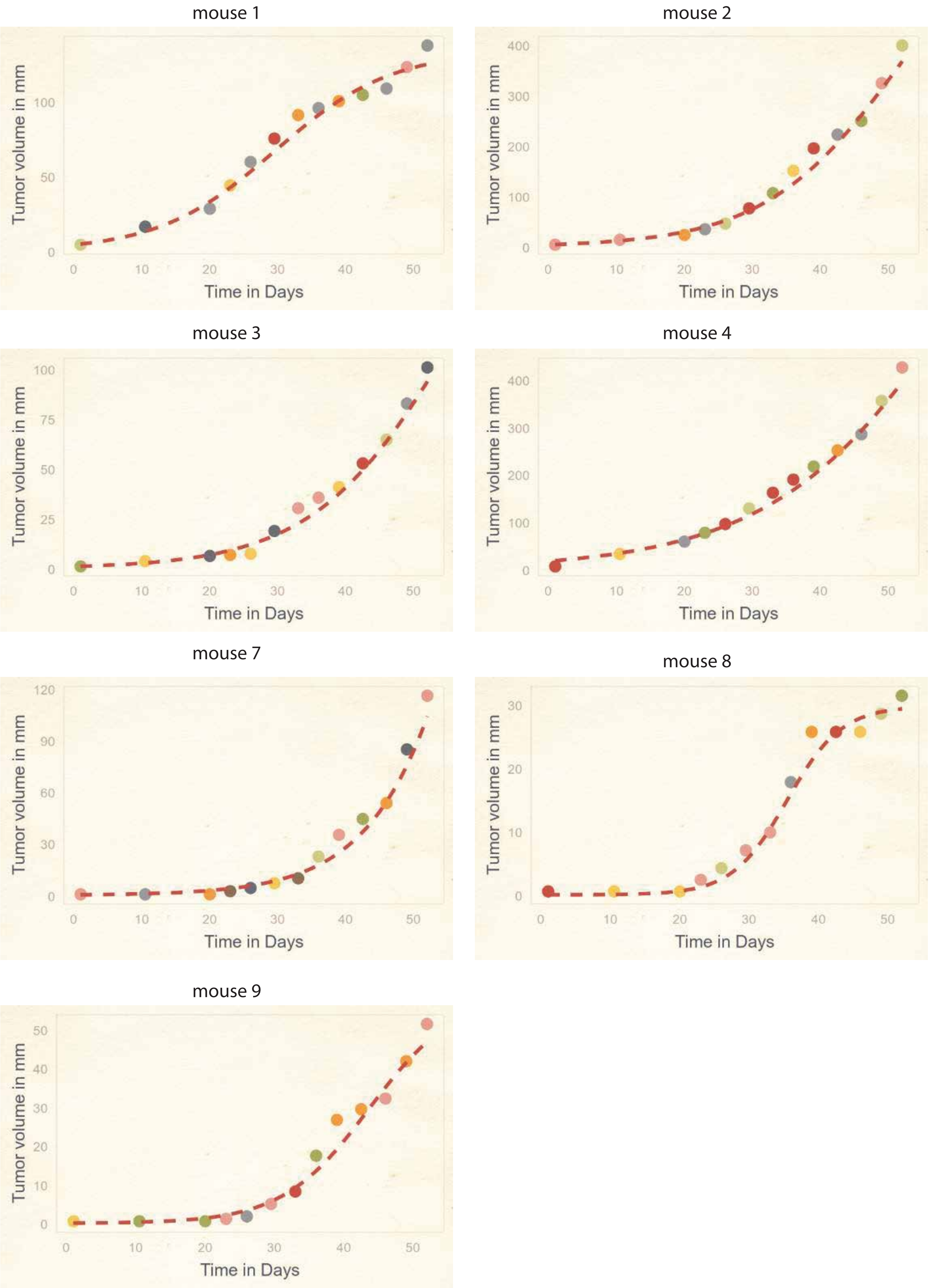

**Supplementary figure 30.** MDA-MB-231 breast cancer cell line

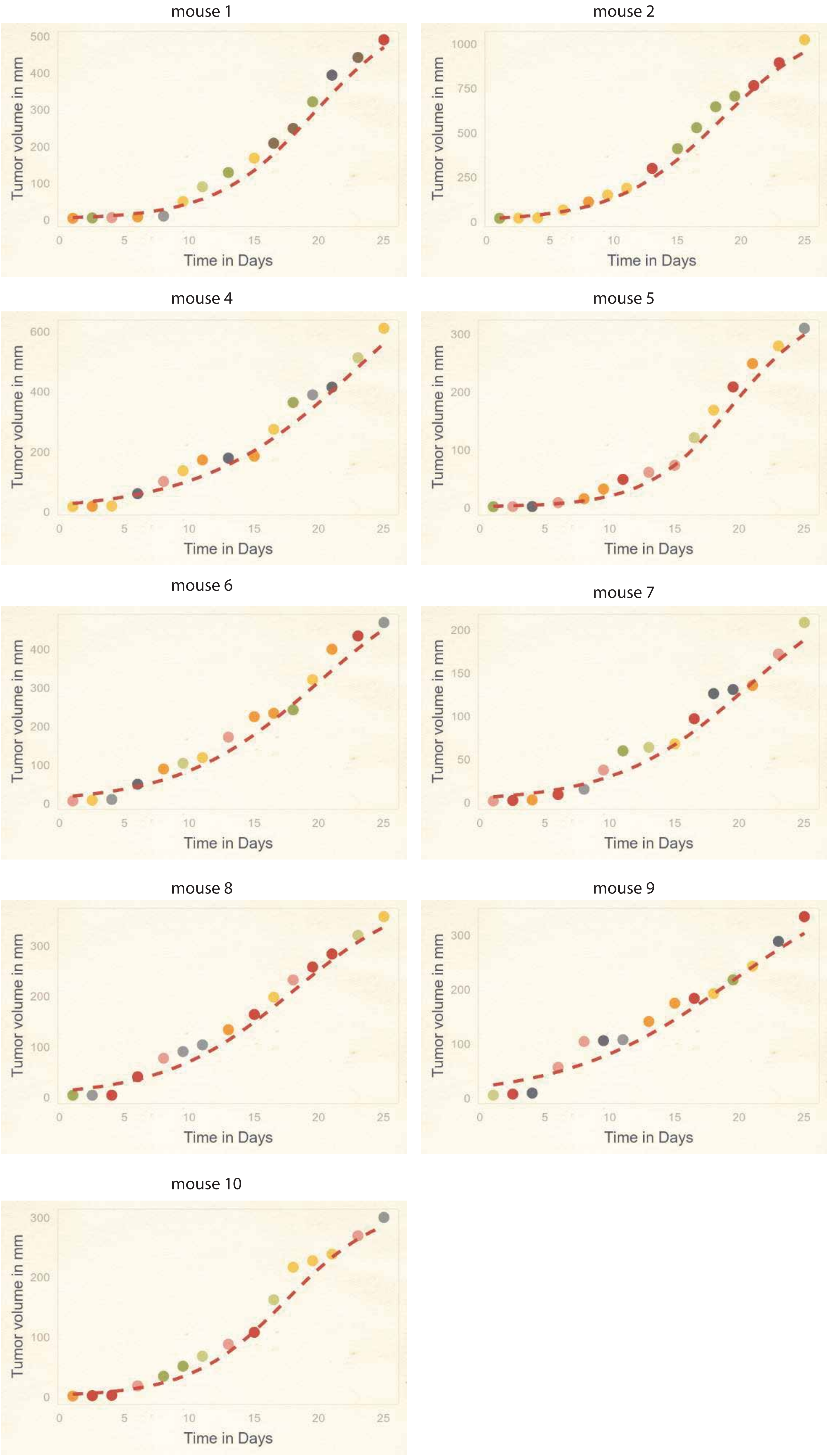
